# Homeostatic feedback model of energy metabolism with adaptive enzyme levels exhibits problem solving behavior

**DOI:** 10.64898/2026.05.07.721661

**Authors:** Axel de Baat, Michael Levin

## Abstract

Metabolic networks are conventionally described as homeostatic systems that stabilize flux, energy charge, redox balance, and metabolite availability under perturbation. Whether the same feedback architectures can also generate learning-like, experience-dependent adaptation has not been systematically tested. Here we construct a coarse-grained dynamical model of mammalian energy metabolism consisting of coupled ordinary differential equations spanning glucose, glutamine, fatty acid, and oxidative phosphorylation pathways, with Michaelis–Menten-type fluxes, product-inhibition feedback, and an adaptive enzyme-capacity layer whose adjustments are debited directly from the ATP pool. We selected parameter sets by survival across a factorial sweep of environmental noise and regulatory cost. This architecture proves sufficient for non-associative learning. Priming with one perturbation alters the response to a second, producing habituation-like behavior when the first perturbation opens an alternative energetic route and sensitization when it shifts load onto the pathway the second perturbation targets. The memory is localized: persistent displacement concentrates in triglyceride and lactate storage buffers as well as NADH and ATP high-degree cofactor nodes, while glucose, pyruvate, NADPH, ROS, glutamine, and free fatty acids return to baseline. Removing the adaptive layer increases metabolite-level displacement, indicating that adaptive regulation does not erase the trace of experience but relocates it from metabolite concentrations to enzyme configuration. Repeated inhibition of glucose import ratchets the system into a persistent high-lactate state with no genetic change, a non-genetic route to a Warburg-like phenotype. The same models reproduce ischemia-reperfusion dynamics, with an ATP nadir followed by a reperfusion ROS spike, although nothing in their selection targeted this behavior. Notably, the controller is greedy and one-step, with no look-ahead, no memory variable, and no representation of its own energetic cost, indicating that dedicated memory machinery is not required for metabolic systems to become experience-dependent. In rigorous habituation experiments this controller proved too powerful to develop habituation however. We therefore tested different controller architectures finding that perfectly adaptive controllers cannot habituate, suggesting the requirement of imperfect controllers, states that are (partially) obscured from the controller, or lack of full settling of the system between perturbations.

## Introduction

Metabolic networks are usually described as homeostatic systems: they regulate flux, preserve energy balance, and buffer perturbations [1, 2]. Recently, a number of unconventional dynamical systems have been shown to exhibit functions that match classic paradigms in behavioral science [3-5]. Learning dynamics in gene-regulatory networks and other molecular pathways have significant implications in biomedical settings [6-10]. An important knowledge gap is whether the feedback architectures present in metabolic networks can support learning-like behavior. Can prior metabolic experience improve a system’s response to future challenges? This question matters because metabolic adaptation contributes to robustness, stress tolerance, and therapeutic resistance, yet many existing models are either too static to capture adaptation or too parameter-rich to make emergent behavior interpretable. Here, we address this gap by developing a coarse-grained dynamical model of mammalian energy metabolism and using it to test whether adaptive enzyme regulation, energetic control costs, repeated perturbation, and environmental variability can produce experience-dependent improvements in metabolic homeostasis.

Metabolism constrains nearly every cellular process by supplying ATP, reducing equivalents, and biosynthetic precursors. A cell that cannot maintain energy charge, redox balance, or metabolite availability will fail to proliferate, migrate, or respond appropriately to signals, regardless of its genetic state [11]. This dependence on metabolic stability is maintained through layered regulatory mechanisms, including allosteric feedback, nutrient transport, post-translational modification, enzyme synthesis and degradation, and transcriptional remodeling. Together, these mechanisms allow cells to stabilize metabolite concentrations while also adapting to changing internal and external conditions.

A striking consequence of this architecture is metabolic plasticity. When a pathway is inhibited by a drug, mutation, or environmental stressor, cells may reroute flux through compensatory pathways, alter nutrient uptake, or change enzyme abundance to restore homeostasis. This adaptive capacity is clinically important because it contributes to resistance: inhibition of one metabolic route can often be bypassed by increased reliance on alternative carbon sources or energy-producing pathways [12]. Such behavior suggests that metabolism is not merely passively robust, but may actively search for alternative solutions that preserve viability.

This possibility connects metabolism to a broader question in biological information processing: whether non-neural biochemical networks can exhibit memory, conditioning, or learning-like dynamics. Prior computational work has shown that gene regulatory and biochemical network models can display forms of memory, Pavlovian-like conditioning, and experience-dependent plasticity [13-15]. These findings motivate the hypothesis that metabolic networks, with their dense feedback loops and adaptive control mechanisms, may also support learning-like behavior. However, this has not been systematically examined in a model designed to expose how metabolic feedback, enzyme adaptation, energetic cost, and environmental variation interact.

Existing metabolic modeling frameworks are not ideally suited to this question. Constraint-based approaches such as flux balance analysis are powerful for analyzing steady-state flux distributions, but they generally do not capture transient adaptive dynamics [16]. Kinetic ODE models can represent time-dependent metabolite concentrations and fluxes, but they often require many poorly constrained kinetic parameters, making interpretation and identifiability difficult [17, 18]. To probe the possible learning-like behavior in metabolic systems, a model that is dynamic enough to capture adaptation but simple enough to make the relationship between structure, parameters, and behavior interpretable is necessary.

We designed a coarse-grained dynamical model of cellular energy metabolism to meet this need. The model represents core mammalian energy metabolism as coupled ordinary differential equations tracking major metabolite pools across glucose metabolism, glutamine metabolism, fatty acid metabolism, and oxidative phosphorylation. Pathway fluxes follow Michaelis–Menten-type kinetics with product-inhibition feedback, so that deviations from homeostatic setpoints alter production and consumption rates. Enzyme levels are represented as adjustable pathway-capacity multipliers, updated by an adaptive controller that tests whether small enzyme changes reduce deviation from metabolic setpoints. Crucially, enzyme adjustment carries an ATP cost, coupling regulation to the metabolic state it controls.

This framework is not intended to reproduce quantitative fluxes in a specific cell type. Instead, it is designed to ask whether a metabolically realistic feedback architecture can generate learning-like adaptation. We use it to test whether prior perturbation improves later recovery, whether adaptive enzyme regulation is required for this effect, and whether environmental variability expands or restricts the repertoire of reachable adaptive states. By combining coarse-grained metabolic structure with explicit dynamics and control costs, the model provides a tractable way to study how homeostatic regulation may give rise to experience-dependent metabolic plasticity.

## Results

### A cost-constrained adaptive enzyme layer navigates conflicting homeostatic constraints

To study homeostatic control via simulations, we constructed an ordinary differential equation model of metabolism that advances intracellular metabolite concentrations forward in time. The model tracks core energy, redox, carbon, and lipid pools (ATP, NADH, NADPH, glucose, pyruvate, lactate, glutamine, ROS, free fatty acids, and triglyceride) relative to predefined realistic setpoints and an explicit extracellular environment (e.g., glucose, glutamine, lactate, oxygen, fatty acids). Fluxes are computed for major processes spanning uptake/transport, central carbon metabolism, redox/ROS handling, and lipid storage/oxidation (e.g., glycolysis/PPP/TCA/ETC, LDH/MCT, ROS scavenging, fatty-acid import, de novo lipogenesis, esterification/lipolysis, and β-oxidation) using standard saturating kinetics and feedback “gates” that suppress or activate pathways when metabolites deviate from their respective targets (Fig. 1A).

**Figure 1.**
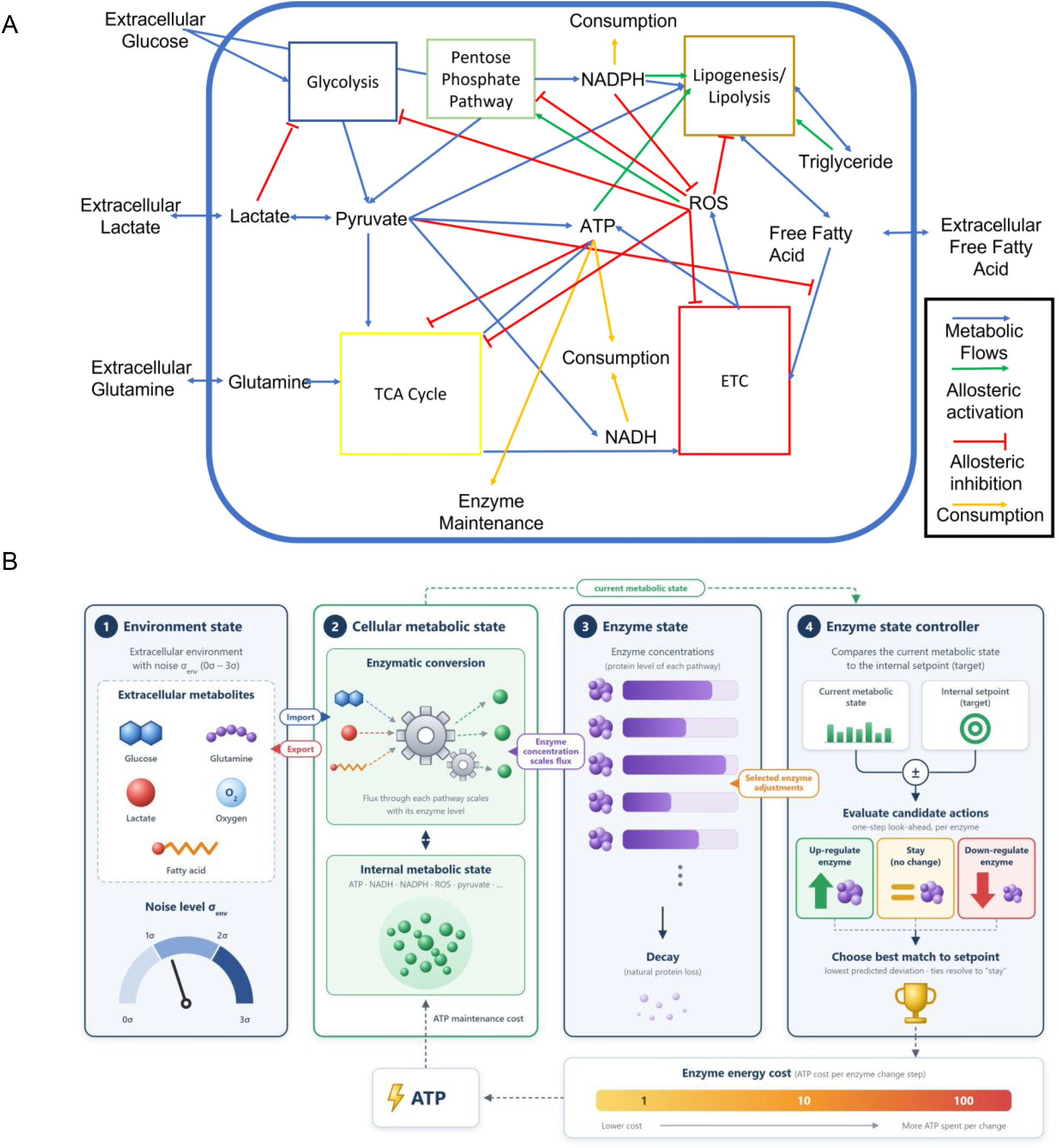
Model structure and representative simulations. (a) Schematic overview of metabolite flows (blue arrows), activating feedbacks (green arrows), consumption (orange arrows), and inhibitory feedback (red lines). (b) Outline of the adaptation process used to adjust internal enzyme levels that scale conversion rates to maintain internal setpoints.

The model additionally features an adaptive “active” enzyme concentration layer: each pathway enzyme is represented as a dimensionless concentration multiplier (initialized to 1.0) that scales selected kinetic parameters (e.g., Vmax terms) via an enzyme-parameter mapping.

The system generates the energy (represented by the cofactors NADH, ATP, and NADPH) that it uses to maintain its enzyme concentrations and support basal consumption representing cellular function. These dynamics were chosen to represent rapid changes in effective enzyme availability through sequestration, post-translational modification, degradation, or enzyme production, together with enzyme-maintenance energy consumption as a proxy for cellular processes (Fig. 1B).

Figure 2 shows representative baseline simulations of a parameter set selected at mild (Env1_EC10) and challenging environmental conditions (Env2_EC100). At initialization, the system explores other many unstable constellations before reaching a more stable state that persists with minimal drift. The model is therefore able to exhibit problem-solving behavior via navigation of conflicting constraints; in this case, changes in enzyme concentrations moving metabolites in orthogonal directions relative to their setpoints.

**Figure 2.**
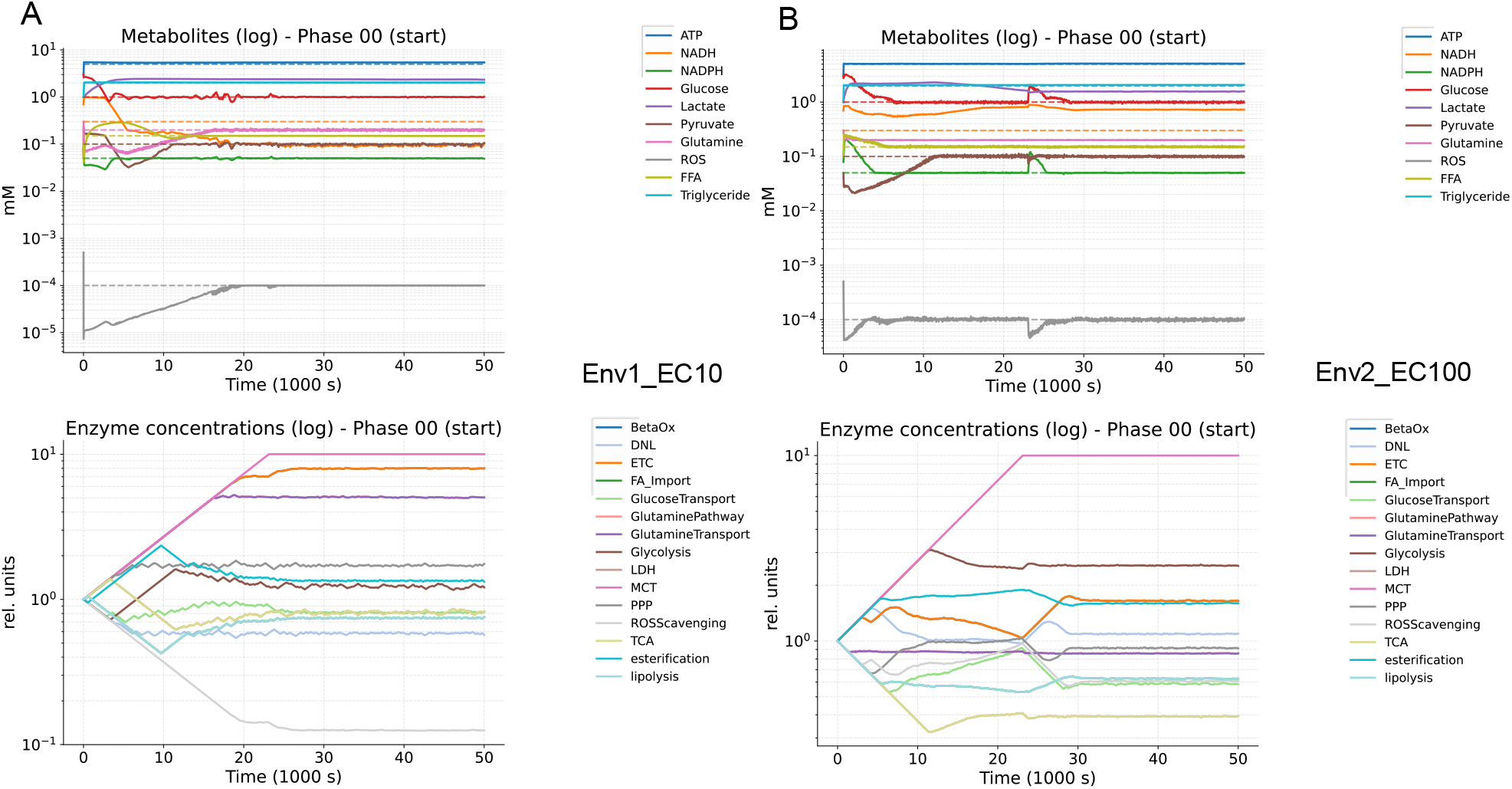
Baseline simulations show models navigating enzyme space to reach a steady state.

### Harsher environments drive deeper parameter-space search and directional shifts in parameters related to transport capacity

Experimentally derived enzyme kinetic parameters show large biases depending on method and experimental conditions and therefore are difficult to harmonize. Additionally, there are no datasets to our knowledge that would capture the richness of dynamics displayed by this model. We therefore chose a nature-inspired simple evolutionary approach, searching for parameter sets that stabilize below a certain stability threshold. Parameter sets were terminated early when any tracked metabolite moved outside either absolute bounds for specified species (e.g., ROS) or fold-bounds around the setpoint using an effective setpoint floor.

Using a fixed seed, we performed deterministic Latin hypercube sampling in log space around the model’s default parameter set.

To mimic existing biological constraints on evolution, candidate parameter sets were further adjusted into a physiologically plausible regime. Each sampled set was passed through a literature-motivated soft-repair step that enforces qualitative flux-capacity relationships before simulation— for example, lactate import capped relative to export (consistent with net export bias under glycolysis), reverse capacities limited relative to forward capacities for reversible pathway modules (glycolysis/PPP/TCA/ETC/glutamine–TCA), PPP not exceeding glycolysis, ETC capacity forced to exceed TCA capacity to avoid chronic NADH accumulation, and substrate import capacities tied to downstream utilization (e.g., glutamine import vs glutamine oxidation capacity, fatty-acid import vs export bias in rich media) (Fig. 3).

**Figure 3.**
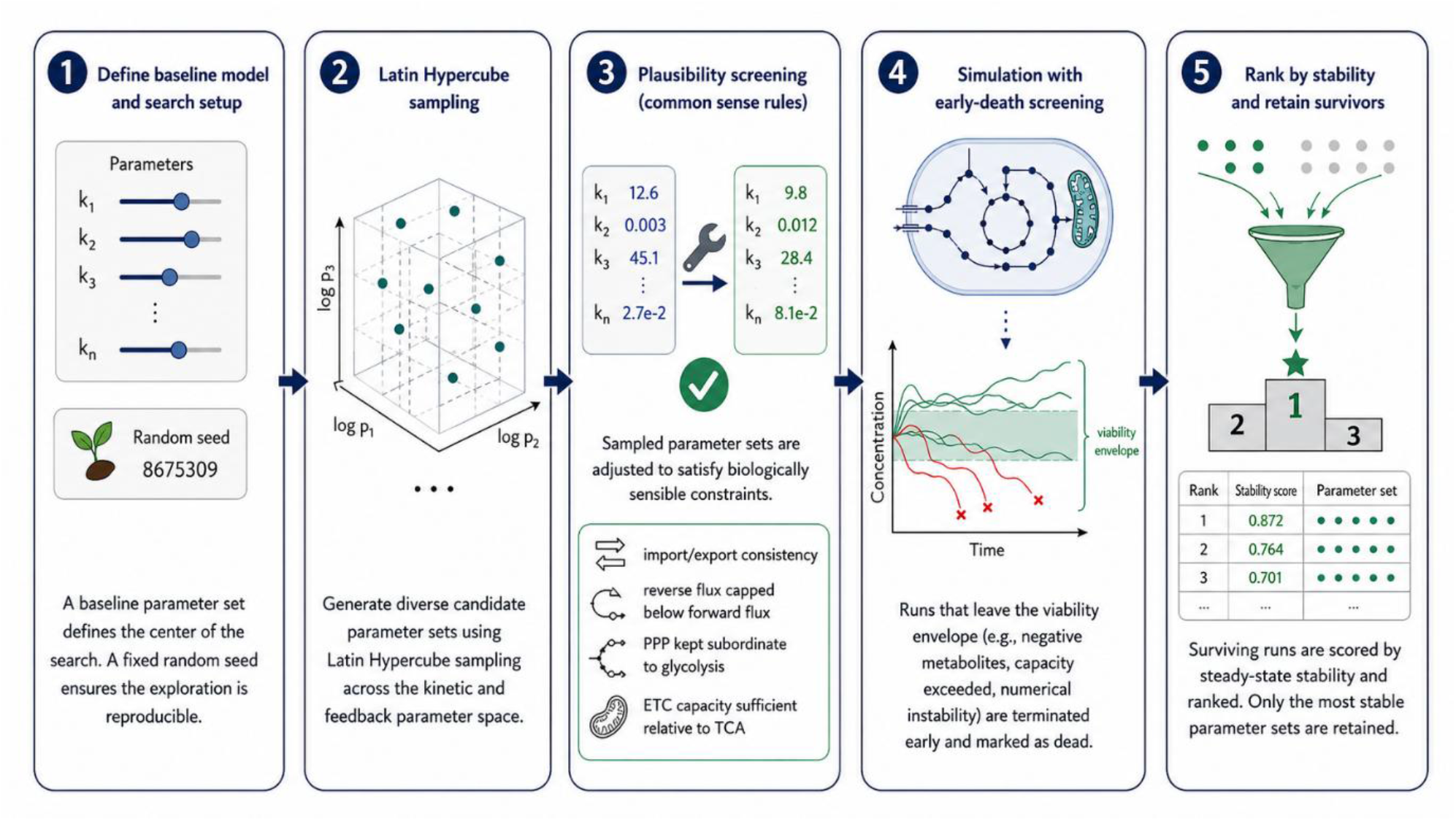
Graphical representation of the parameter search process.

To test how internal energetic demands and environmental noise shaped parameter-set selection, we evaluated the same sequence of sampled parameter sets under each condition. We varied mean-preserving multiplicative log-normal noise at each internal sub-step (Env; 1-3 sigma, represented as Noise) to probe the hedging-versus-efficiency tradeoff and varied the energetic costs (EC) associated with increasing enzyme levels (1-100-fold baseline, represented as EC#).

We then investigated the extent to which the simulation environment shapes the selected parameter values in the surviving sets. A more challenging environment results in deeper exploration of parameter space as evidenced by the number of parameter sets tested before a viable parameter set is reached (Fig. 4A). We then compared the selected parameter values per cohort to a randomly selected population. This gives a readout of what parameters have a large impact on system survival and are therefore constrained by selection. Parameters governing metabolite uptake and maximum reaction rates are tightly constrained by selection (Fig. 4B). We also evaluated whether the mean shifted lower or higher relative to a randomly selected population (Fig. 4C). *Km_lactate_export*’s mean shift shows this ascending pattern with increasing environmental difficulty, likely reflecting a suppression of lactate export to increase glucose efficiency. *Vmax_glucose_import* shows the reverse pattern, in line with adaptation to increasing energy demands.

**Figure 4.**
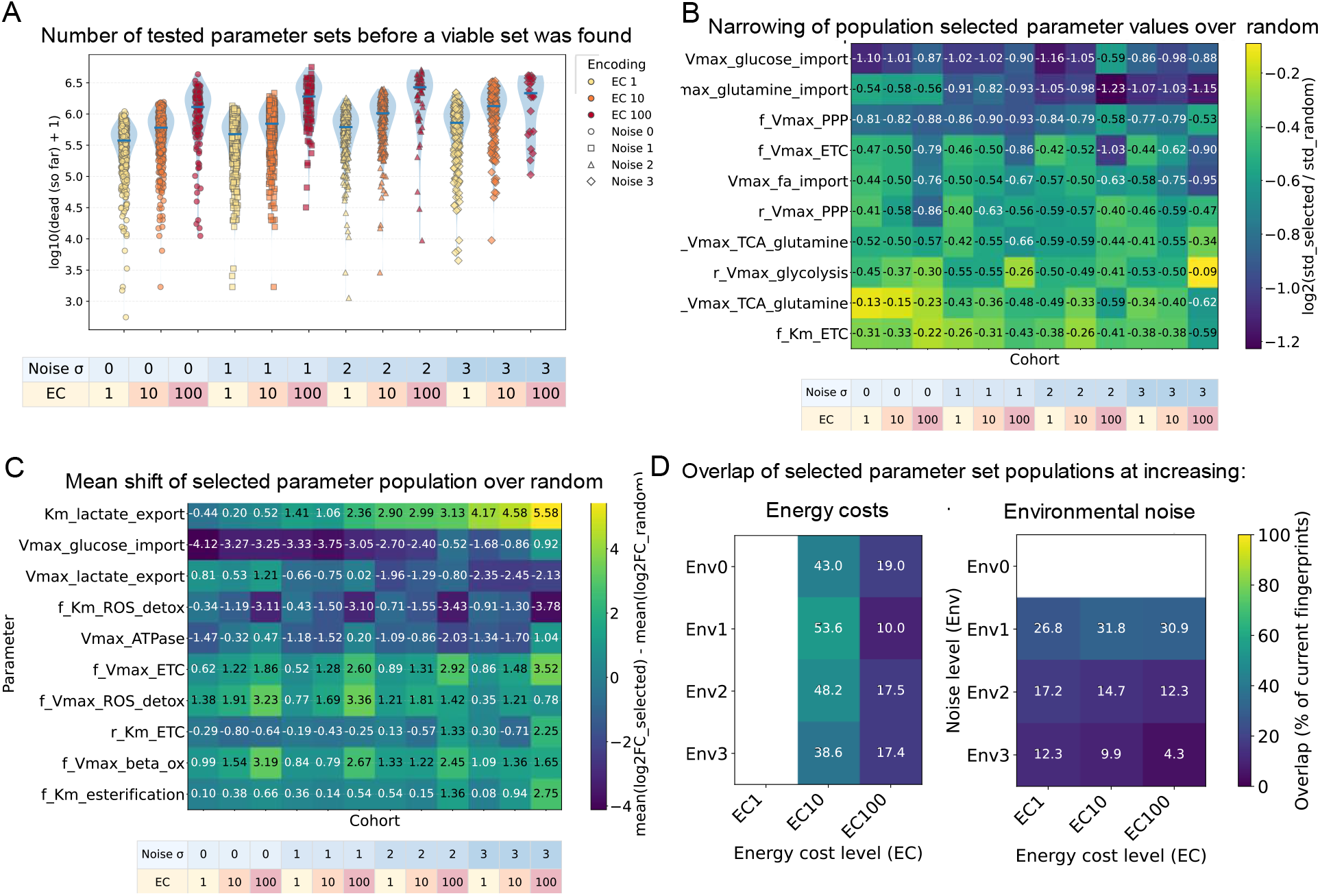
The simulation environment shapes parameter-set selection and the individual parameter values retained. (A) Number of tested parameter sets before a viable parameter set is found; higher values indicate deeper exploration of parameter space, (B) Contraction of selected parameter values compared to a randomly selected population, (C) Mean shift of parameter values compared to a randomly selected population, (D) Overlap between parameter-set populations with increasing energy costs (left) or environmental noise (right).

All parameter search simulations were sampled from the same randomly generated sequence of parameter sets, providing insight into the extent to which environmental conditions affect what parameter sets are selected. Overlap was compared in surviving parameter sets between simulations where Gaussian noise was applied to the environment (Env#) and where the energy costs associated with changing enzyme concentrations were scaled (EC#) (Fig. 4D). More overlap was observed between populations sampled at increasing energetic costs at every noise level than vice versa; however, this is likely because a transition from no noise to noise is more impactful than just scaling up the energy costs. When comparisons were made starting from Env1, the impacts are largely similar. More stringent simulation conditions therefore result in deeper exploration of the parameter space, analogous to how extreme conditions result in extreme biological characteristics due to lack of rapid niche filling, allowing for a deeper search of biological potential leading to the evolution of extremophiles.

### Intermediate environmental variability selects for efficient, generalist metabolism; extreme variability selects for high-throughput

We then investigated whether differences in parameter values manifest as functional differences in metabolism by profiling the metabolic fluxes of the different simulations. Each parameter set was simulated long enough to reach a non-equilibrium steady state, and the final 10,000 steps (1,000 seconds) were considered for analysis. For every parameter set, the model’s stepwise pathway fluxes were recorded, and the mean net rate for each pathway was computed over the final simulation window. Net rates were defined as the difference of forward and reverse fluxes for reversible pathways (e.g., glycolysis, PPP, TCA, glutamine-to-TCA) and as import minus export for transport processes. Total flux (total pathway activity) was defined as the sum of absolute values of these net pathway rates across the set of pathways considered (e.g., glycolysis, PPP, TCA, LDH, MCT, ETC, and DNL/ROS), yielding a single scalar per parameter set that reflects overall metabolic throughput.

The main factor influencing the total flux is the energy costs of the system, which synergizes with environmental noise, mostly in the extreme noise conditions (Fig. 5A). The median ETC rate of the window shows that this increase in total flux is driven in large part by ETC activity (Fig. 5B), which reflects the increased energy demand as ETC is the most efficient means of generating ATP required for modulating enzyme concentrations. A big driver ATP consumption is the maintenance of the enzyme concentrations (Fig. 5C), EC10 seems to most closely related to estimated proteome maintenance estimations of ∼20% [19]. The increased glucose uptake rate is also driven by the increased need for NADPH generation to detoxify the ROS generated by the ETC which is increased for efficient energy production (Fig. 5D).

**Figure 5.**
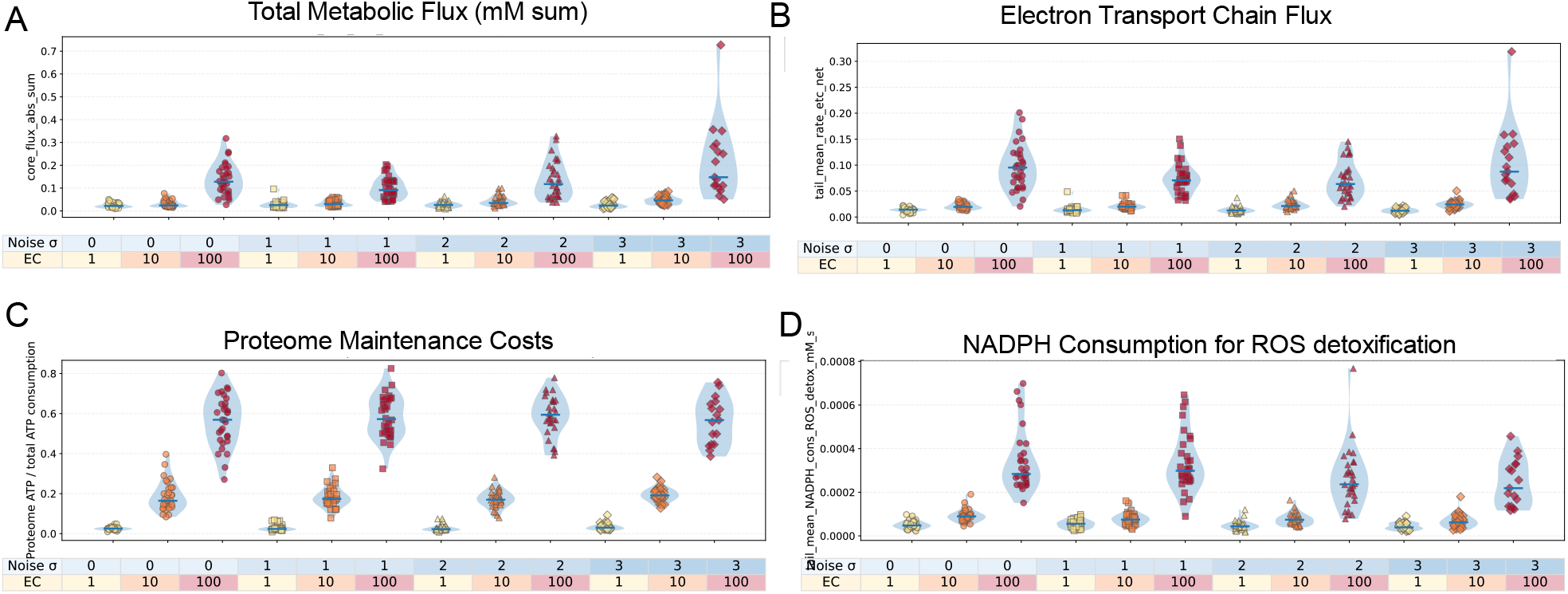
More challenging environmental conditions result in more energy-intensive metabolism driven by the electron transport chain. (A) The sum of all the net fluxes, (B) Mean electron transport chain flux, (C) ATP consumption associated with maintaining enzyme concentrations normalized by total ATP consumption, (D) NADPH consumption associated with the detoxification of reactive oxygen species for all the parameter sets at steady state.

When carbon sources are compared by uptake for all the selected parameter sets, a high diversity in dominant carbon sources is observed. With increasing noise, we see a shift away from glutamine metabolism and especially at high energetic costs, an increase in glucose-dominant metabolic states, away from fatty-acid-dominated metabolism, likely due to the increased ROS burden of lipolysis (Fig. 6, left). The source of ATP production confirms this by showing a clear shift towards the most efficient oxidative metabolism with increasing energetic demands, where environmental noise selects for a small population of mixed metabolism, which is negated by extreme energetic demands (Fig. 6, right). Notably, low energetic demands result in selection of glutamine dominant sets, as glutamine is the most predictable, but low energy yielding, fuel.

**Figure 6.**
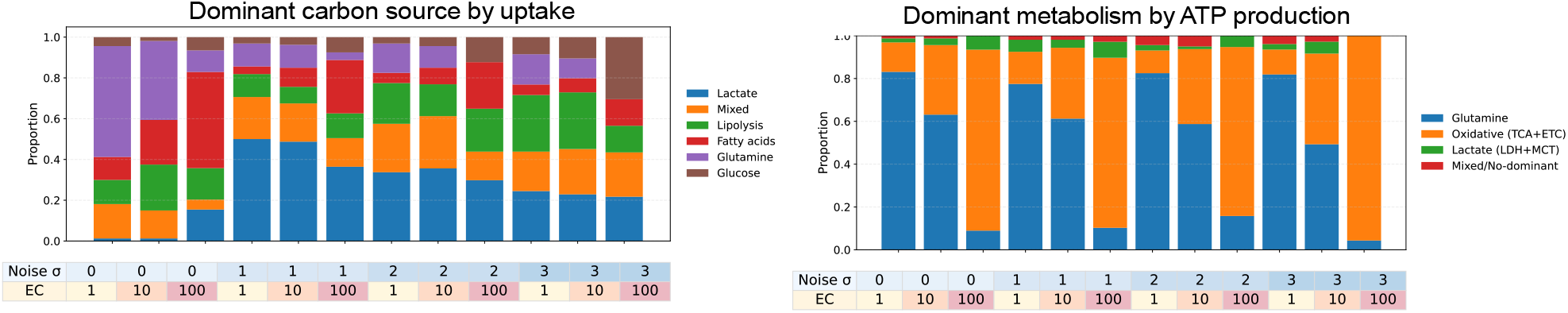
Metabolic characteristics of the different cohorts. Dominant carbon source according to transport and conversion (Left). Dominant metabolism according to ATP yield (Right). Env# denotes the noise level in standard deviations, and EC# denotes the energy-cost factor associated with changing enzyme concentrations.

### Ischemia–reperfusion dynamics emerge without specific selection

To probe whether the system could exhibit realistic, disease-relevant behavior, we created a simulation for ischemia-reperfusion (I/R). During ischemia, circulation is restricted, resulting in a drop in oxygen tension and metabolite supply. This causes respiration and ATP to decline; in response, the metabolic system adjusts to maintain homeostasis. When the blockage is resolved and the tissue is re-perfused, the system has adapted to the new environment, and this rapid change back to oxygen-rich conditions results in a deleterious spike in reactive oxygen species. In the simulation, an ischemic event is represented as a reduction of oxygen and glucose in the environment to 0.5% and approximately 4.5% of their baselines, respectively. The left panel of Figure 7 features a representative simulation highlighting the ATP nadir and subsequent spike in reactive oxygen species. The scatter on the right side of Figure 7 plots the ROS spike ratio against the ATP nadir, showing this behavior is present in varying degrees in all selected parameter sets, suggesting this behavior is an intrinsic property of the model structure.

**Figure 7.**
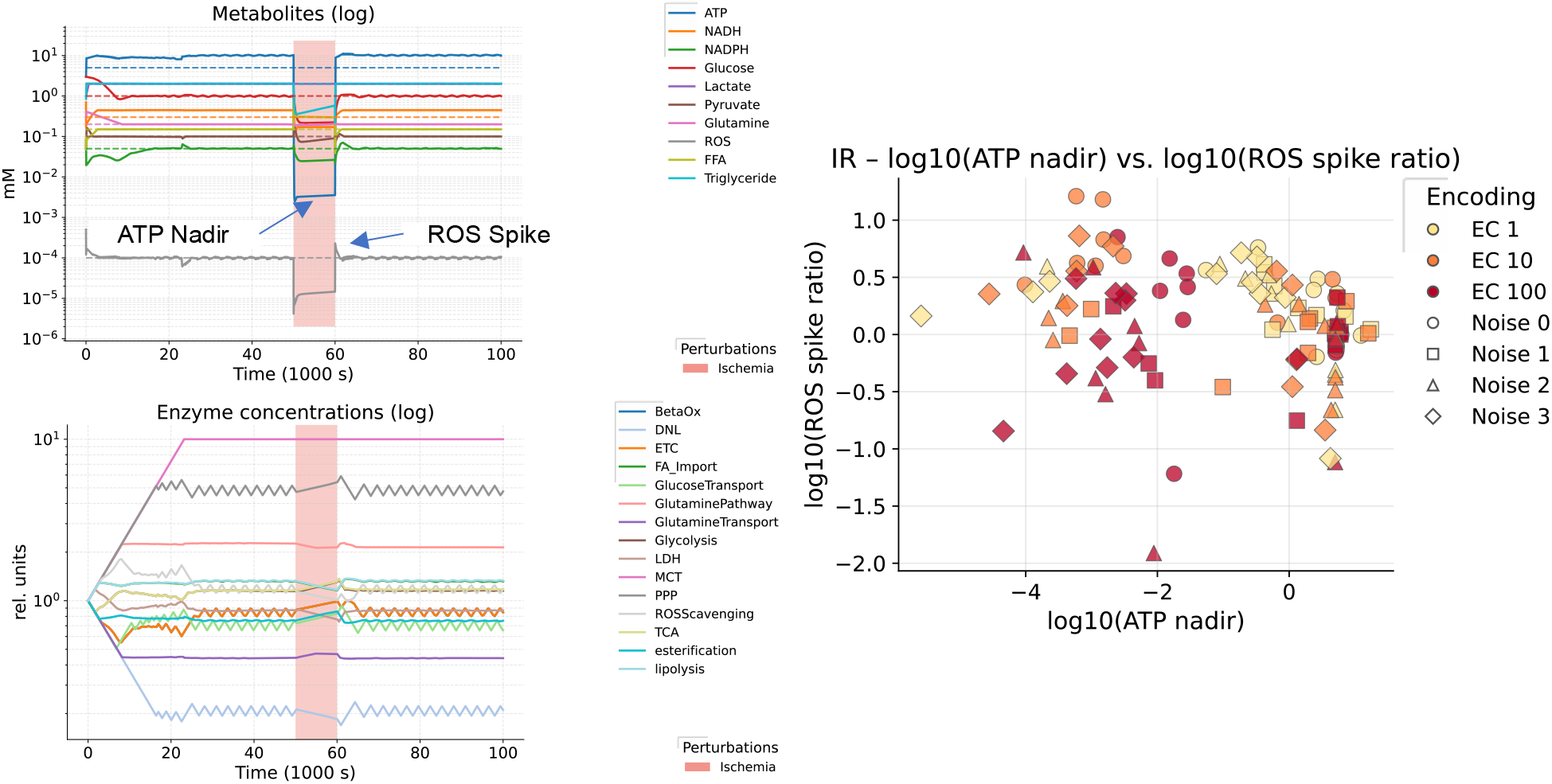
Representative ischemia-reperfusion simulation (left). Dashed lines indicate setpoints. ROS spike ratio, indicating the increase in ROS, relative to the ischemic phase, after oxygen and glucose restoration, plotted against ATP nadir, defined as the lowest ATP concentration (mM) during oxygen and glucose deprivation (Right).7

### Persistent metabolic memory is localized to storage buffers and cofactor hubs

To further probe the homeostatic capabilities of the systems, we performed simulations in which the system was perturbed to study its recovery. The following types of perturbations at two strengths were included: changing the environment (change_env) where extracellular environmental metabolites (e.g., lactate and glucose) are reduced, inhibitors that inhibit key metabolic reactions, and increasing the baseline consumption of certain cofactors thereby increasing metabolic burden, mimicking an insult leading to a metabolically taxing stress response. The top ten parameter sets selected based on stability were run until they reached a NESS; subsequently, all candidate single-step perturbations were exhaustively evaluated on this NESS. Each candidate was tested under a pulse regime (apply for a fixed 10,000 s perturbation window, then remove the perturbation while retaining the current metabolite state and enzyme concentrations) at low and high strengths of 0.3 and 0.7, corresponding to a 30% and 70% decrease in environmental metabolites and enzyme activity for inhibitors, and a 30% or 70% increase in consumption rate for consumption-increase perturbations. The parameter sets were then scored by the persistent metabolite displacement (d_met) relative to the pre-perturbation steady state after the 40,000-s recovery window. The highest-ranked perturbation by d_met was used as the first step in a program and used as the baseline for the subsequent round, yielding the most effective two-step protocol tailored for every parameter set (Fig. 8A).

**Figure 8.**
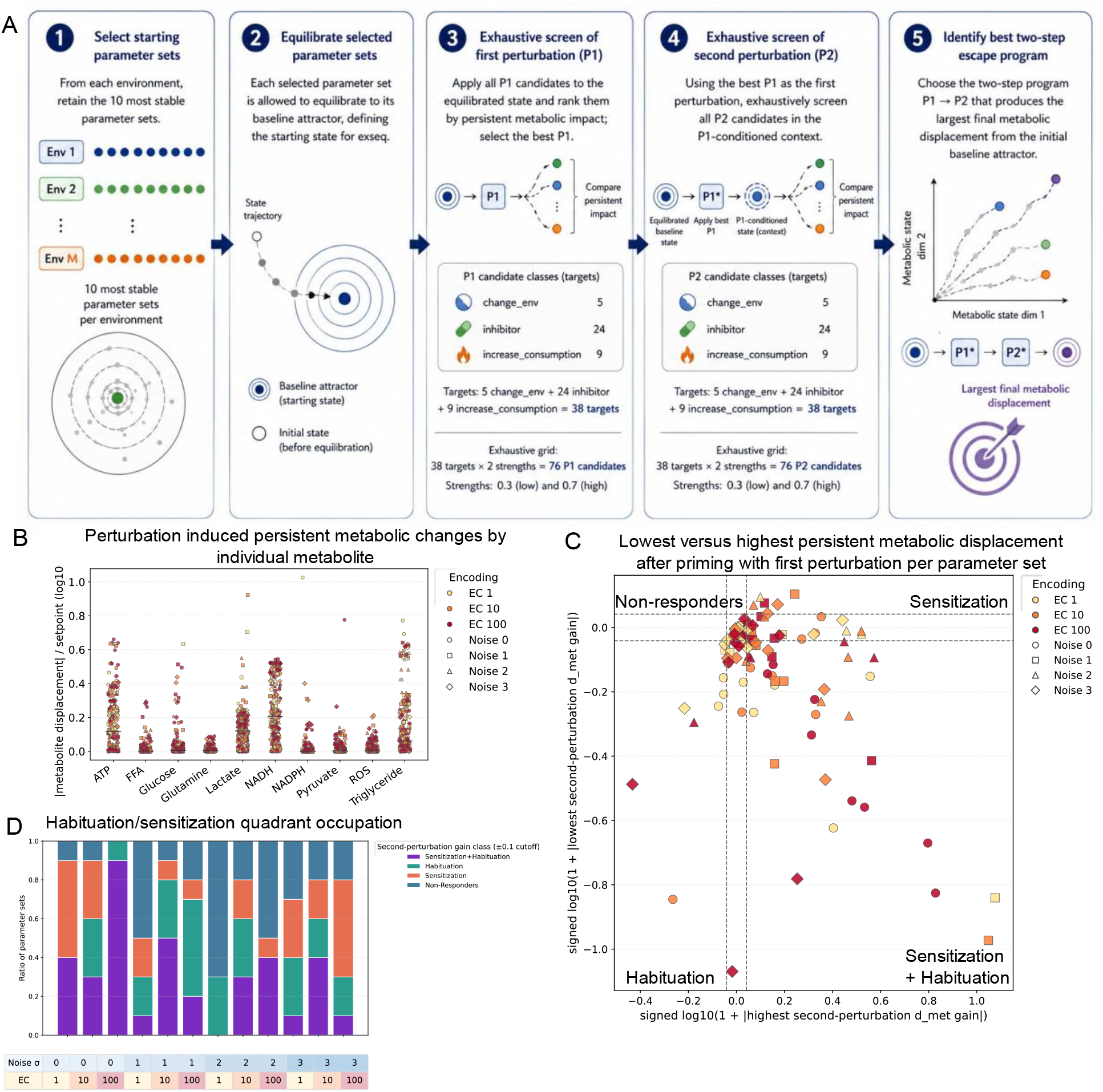
Exhaustive perturbation screening identifies sensitizing and habituating behavior localized to specific nodes in the metabolic network. (a) Graphical overview of the perturbation screening process, (b) Perturbation-induced metabolic changes per metabolite normalized by its respective setpoint, (c) Scatter showing minimal second-perturbation-induced displacement plotted against the maximal displacement for every selected parameter set, (d) Subdivision by environment of quadrant occupation, corresponding to habituating and sensitizing behavior.

**Figure 9.**
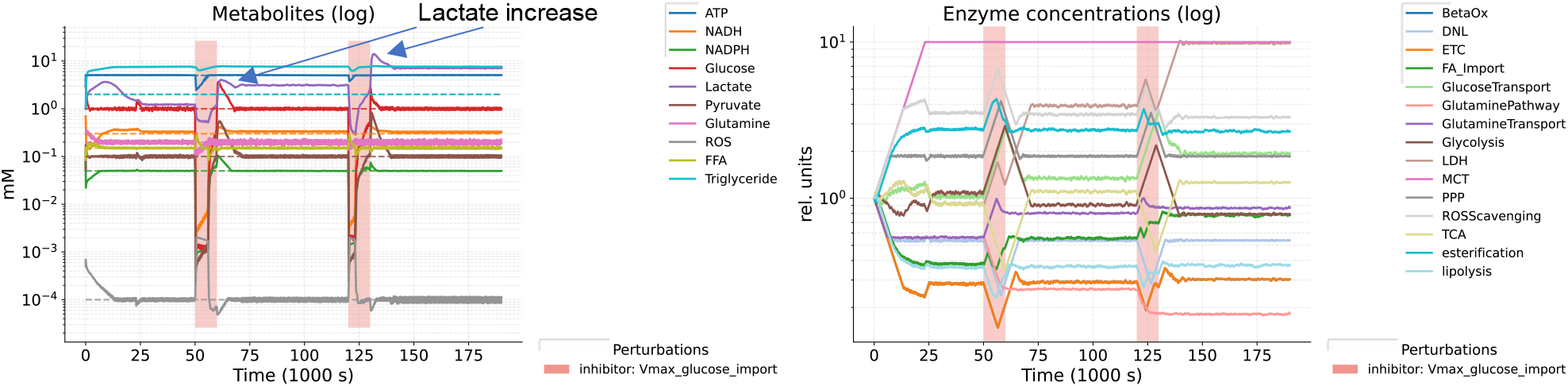
Repeated glucose import inhibition produces a Warburg-like increase in lactate.

To investigate whether specific metabolites were more amenable to persistent metabolic changes, we normalized each metabolite’s persistent displacement in response to every perturbation by its respective setpoint (Fig. 8B). Specific nodes in the metabolic network (ATP, lactate, NADH, and triglyceride) show a higher propensity for being displaced by perturbations.

### Priming produces habituation or sensitization depending on whether it routes flux toward or away from the second target

The two-step perturbation protocol allows us to evaluate whether there is a diminished or increased response to the second perturbation, which is indicative of habituation and sensitization, respectively. We therefore plotted the smallest d_met response to the second perturbation against the corresponding largest response for every parameter set (Fig. 8C). We subdivided the scatter into quadrants corresponding to habituating and sensitizing behavior with a 0.1 d_met cutoff for non-responders (dashed lines). Habituation and sensitization behaviors were observed in all environments (Fig. 8D). Endpoints therefore depend on the order of perturbations, not only their identity, which is the operational definition of a system with memory.

### Repeated glucose-import inhibition ratchets the system into a persistent high-lactate state

As an example, one of the most impactful programs was repeated inhibition of glucose import. This resulted in progressively higher lactate levels with each pulse. Here we see a rapid drop in metabolites once glucose import is limited, resulting in rapid changes in enzyme concentrations to return to homeostasis. When glucose import is restored, the system cannot return to its original metabolic state but instead returns to a state with higher lactate. When lactate export is subsequently inhibited, lactate levels increase sharply; as a compensatory response at the enzyme level, glucose metabolism and lactate production are decreased. When the inhibition is removed, the system converges again to a different NESS with higher lactate levels. This represents a non-genetic route to Warburg-like metabolism.

The left panel of Figure 10 is a representative simulation of a habituation-like response. Two pulses of lactate dehydrogenase inhibition (targeting the enzyme that governs conversion between pyruvate and lactate) result in a smaller glucose spike upon the second perturbation. This is achieved by a shift towards lipid metabolism, alleviating the pressure from lactate metabolism by offering an alternative energetic route. The right panel features a simulation characteristic of sensitization. The first perturbation most notably moves the system away from fatty acid uptake by limiting the supply of fatty acids from the environment. When the system is then confronted with inhibition of glucose import, to which it has rerouted part of its metabolism, the system shows a large metabolic displacement and a move back towards fatty acid metabolism.

**Figure 10.**
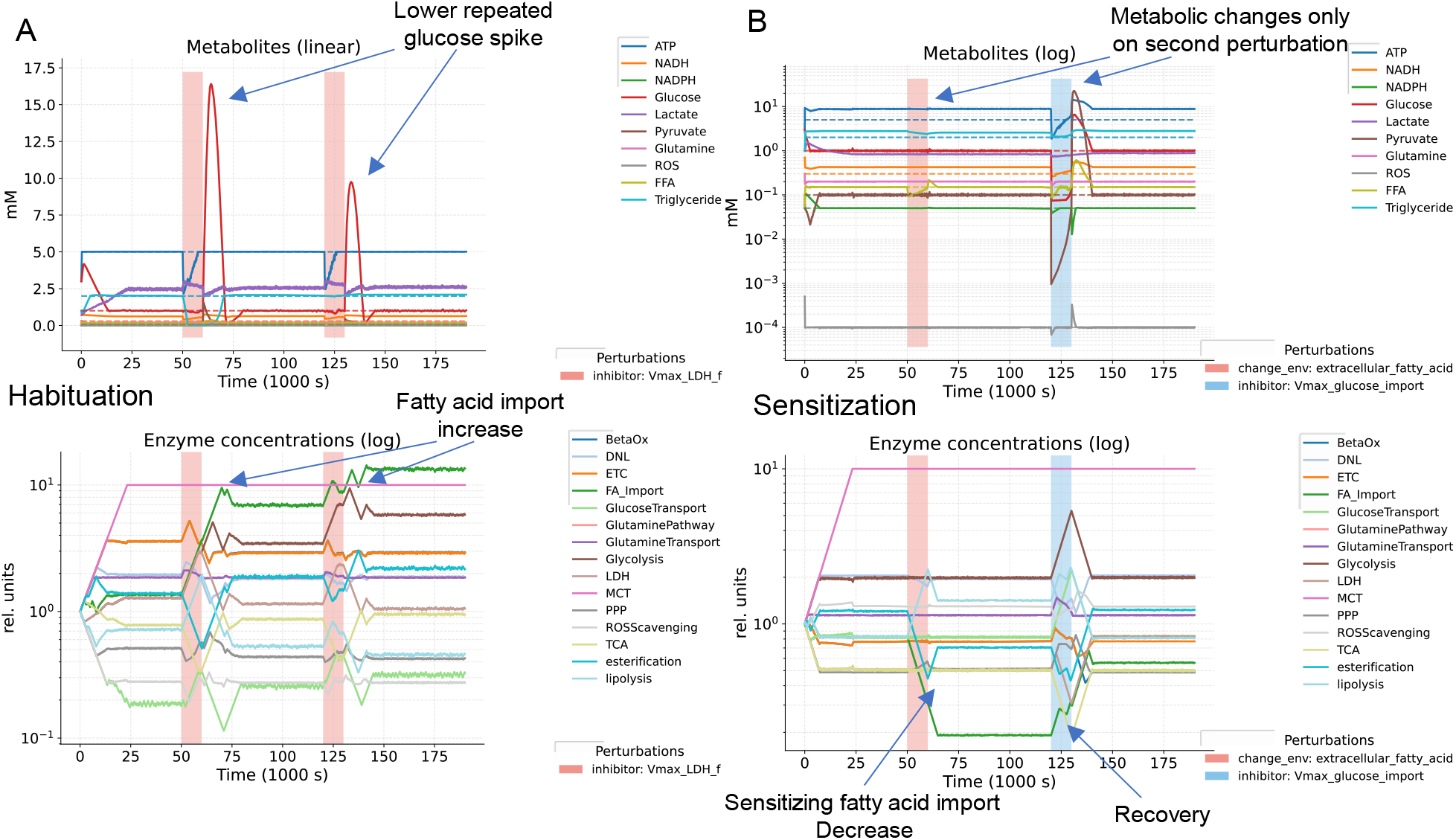
Example simulations of habituating (a) and sensitizing (b) perturbation programs.

### The adaptive layer relocates memory from metabolite concentrations to enzyme configuration

To test how the presence of the adaptive enzyme system affects the dynamics of the model, an additional cohort of parameter sets was selected as before, but without an adaptive enzyme system, relying on the enzyme kinetics and metabolic feedbacks alone. The parameter sets were then subjected to the same exhaustive perturbation profiling as before. The non-adaptive systems show a more pronounced response at the metabolite level to perturbations, compared with systems that have an adaptive enzyme system (Fig. 11A,B). The representative simulations show the robustness conferred by the adaptive enzyme system, whereas the non-adaptive system is unable to recover after perturbations and the perturbations result in large metabolic changes. When we compared pair-ranked programs with the highest metabolic displacement between adaptive and non-adaptive systems, we found that the optimal programs for non-adaptive systems produced a persistent metabolic displacement that was, on average, approximately 10 d_met higher (Fig. 11C).

**Figure 11.**
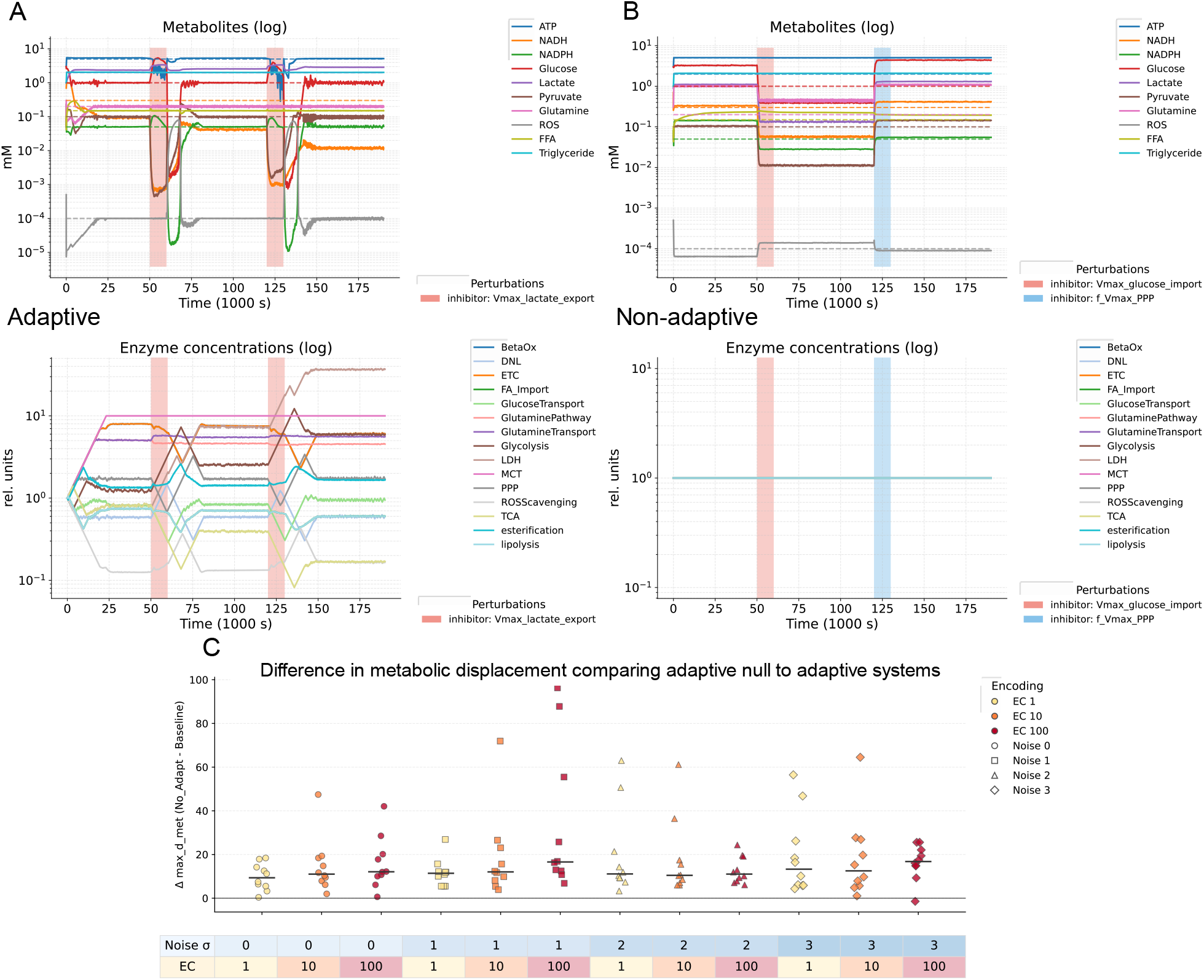
Comparison of models with and without adaptive systems. Representative simulations comparing models with (a) and without (b) an adaptive enzyme system. (c) Difference in metabolic displacement with the optimal program between paired adaptive and non-adaptive parameter sets ranked for stability.

### Rigorous habituation testing shows that the adaptive enzyme system does not habituate when given relaxation, suggesting necessity for incomplete adaptation in habituation

To more rigorously test habituation behavior, we set up a classical habituation simulation with repeated perturbations, a subsequent test with an unspecific perturbation, and ultimately a re-test to see if the habituation relaxes over time (Fig. 12A). In habituation, these repeated measures should result in increasingly smaller responses in both the enzyme and metabolic system (Fig. 12B &C). The basic flat controller does not show habituation behavior in this setup, because the controller perfectly adjusts back and forth in response to the perturbation if given enough time to relax. We therefore tested different enzyme adaptation dynamics to see if the presence of these mechanisms in addition to the standard enzyme controller promote habituation behavior. The tested strategies are summarized in (Fig. 12E). We have tried a shifting setpoint that slowly adjusts to the current metabolic state, in our experiment we found this functioned as a ratchet and moved the metabolic state to unrealistic levels. We tested a modified version that moves the setpoint faster away from its base and with slower movement back and a baseline decay rate back to the base setpoint, this however resulted in too much drift to properly assess habituation. We then tested a mechanism that accumulates a bias on each enzyme’s starting position that is not seen by the controller based on previous peaks. This mechanism did not reduce the peak size, but did improve recovery time, and was therefore not classified as habituating in our framework. We then focused on the interface between the environment and the cell, via modifying the reactions governing metabolite transport. We first tried transport Km desensitization, where the Km (substrate concentration at which the reaction occurs at ½ Vmax) is increased with environmental deviation. This resulted in sensitization however, because the decreased Km was adapted for during rest periods, resulting in more transport capacity which yields higher spikes upon perturbation. As a positive control, we then implemented a controller where the perturbation induced metabolic displacement perceived by the controller is diminished with every perturbation. This results in observable habituating behavior. In order to see if we can encode these dynamics in a more emergent manner, we tried to a mechanism of transporter internalization. This approach was again compensated for by the enzyme system, suggesting that habituation requires either inefficiencies in the controller in reaching its desired state, or states the controller cannot perceive.

**Figure 12.**
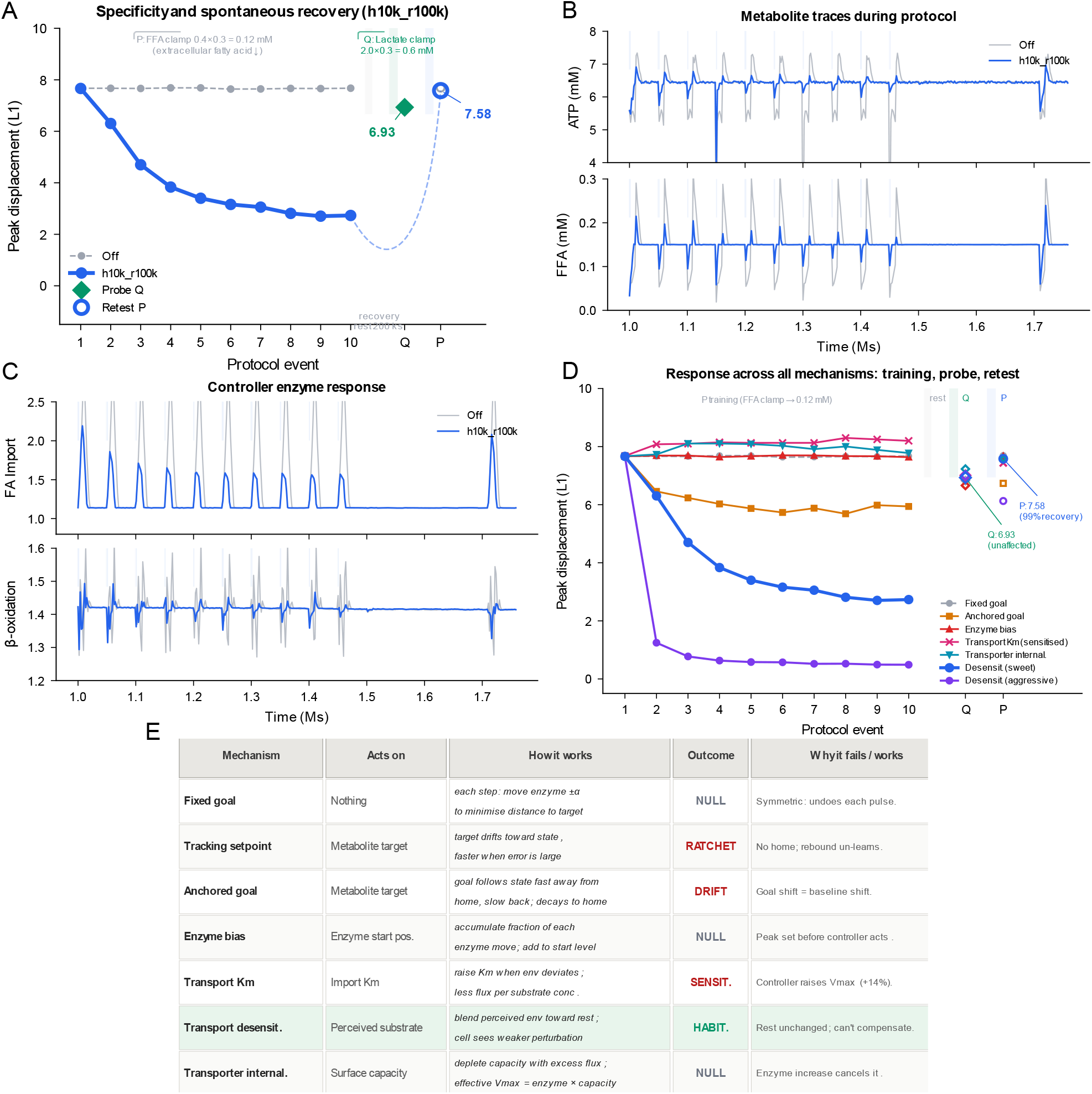
Habituation profiling with alternative controller architectures. (a) Schematic of the classical habituation protocol, (b) expected enzyme response pattern, (c) expected metabolic response pattern, (d) representative simulation results, (e) summary of tested controller mechanisms and their outcomes.

## Discussion

This model captures the emergent behavior of biological systems navigating constraints composed of many, often conflicting, homeostatic feedbacks. This has been performed in a minimal architecture with no look-ahead, no memory variable, no reward signal beyond instantaneous setpoint deviation, no representation of its own cost. We believe these dynamics are fundamental to biology and contribute to habituation and problem solving behavior in the metabolic system.

The simple time-based parameter selection approach, based on survival by maintaining homeostasis. In this study, the environment was reflected in its constraining effect on acceptable parameter values to a narrower bound and or move it away from its ancestral values (in this case the base parameters used as a starting point to sample new parameters). Additionally, we relate the parameter selection to the functional outputs of the system via preferential selection of more energy efficient pathways in environments that are more energetically demanding via a more energy demanding environment. A less energy demanding environment resulted in a preference for glutamine based ATP generation, which is consistent with the observation that *in vitro* cell culture relies more on glutamine metabolism[20]. Another interesting analogous process is cancer cachexia, where the muscles lose glutamine via their constrained metabolism that is used by the cancer that is unconstrained and select for the easiest energy source[21].

The time based component to the parameter selection highlights a mechanism through which extremophiles emerge. In more extreme environments, less parameters are viable and generally die faster, resulting in deeper exploration of parameter space. This could be considered analogous to the emergence of extreme characteristics in organisms evolving in challenging environments via lower niche occupation.

The systems were then screened for perturbations, in order to find ways to elicit learning-like behavior. We then formulated optimal programs that maximally induced persistent changes on a metabolic level. We found an interesting pattern related to what metabolites are more prone to be persistently changed after a perturbation. Metabolites such as FFA, glucose, pyruvate, NADPH, ROS and glutamine are less prone to permanent displacement, whereas NADH, ATP, triglycerides and lactate are, likely due to their respective places in the metabolic network. Triglycerides are for energy storage, which can be used as a buffer to absorb metabolic shocks, Lactate can function as a glucose sink. The energetic co-factors are likely are highly connected nodes in the network.

In some simulations we found Warburg-like metabolism inducing qualities in the programs, suggesting these models are able to show emergent, disease relevant behavior. In others, systems showed a diminished response to the second perturbation after being primed by the first perturbation, a sign of habituation. Sensitization was also found, where priming with the first perturbation increased the impact of a subsequent perturbation. This is striking as these models have a minimal architecture with no look-ahead, no memory variable, no reward signal beyond instantaneous setpoint deviation, and no representation of its own cost.

We then compared the same model framework with and without adaptive enzyme system to understand the differences in dynamics. The null-adaptive model showed more pronounced persistent metabolic changes in response to perturbations, whereas the adaptive model mitigated many of the changes on the enzyme level.

When subjecting the parameter sets to a more rigorous habituation profiling, we found that, when given enough time to relax, our basic flat controller does not produce habituating behavior, suggesting that not allowing the system to fully settle can be a requirement for reaching other equilibria, a necessity for habituation. We then unsuccessfully tested different controllers to see if we can elicit habituation, suggesting that it requires an optimal combination of flexibility and robustness. The only mechanism that did produce habituation relied on partly obscuring the environment the controller sees, suggesting controller inefficiencies are required for habituation as a perfectly robust controller would always revert back to the same setpoint.

This model attempts to capture dynamics at a timescale and resolution not yet reachable by experimental methods. Our attempt has been to highlight the importance of homeostasis as a guiding principle for biological systems and an attempt at generating a model based on qualitative experimentally validated knowledge on the structure of metabolic pathways, allosteric feedbacks and the way in which metabolism maintains homeostasis by modulating metabolic fluxes via enzyme availability on different scales such as production, degradation, sequestration and allosterism [22]. We envision models of this kind to ultimately serve as an interpretable layer that captures qualitative knowledge we have of biological systems that can serve as a means of hypothesis generation, effectively informing experimental design.

## Acknowledgements

This work was supported by the Gates Foundation INV-068658. The conclusions and opinions expressed in this work are those of the author(s) alone and shall not be attributed to the Foundation. Under the grant conditions of the Foundation, a Creative Commons Attribution 4.0 License has already been assigned to the Author Accepted Manuscript version that might arise from this submission. Please note works submitted as a preprint have not undergone a peer review process. The funder played no role in study design, data collection, analysis and interpretation of data, or the writing of this manuscript. We thank Ben Hartl for his feedback on the manuscript.

## Author Contributions

AdB conceived the study, developed the model, performed simulations and analysis, and wrote the manuscript. ML supervised the study and revised the manuscript. All authors read and approved the final manuscript.

## Methods

### Metabolic model

#### Model scope and state representation

We implemented a continuous-state metabolic dynamical model evaluated at discrete time intervals.

The intracellular metabolic state was represented by

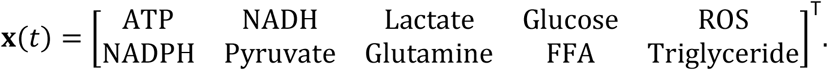

The adaptive enzyme-capacity state was represented by

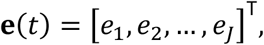

where each *e*_*j*_ is a dimensionless multiplier of the maximum capacities assigned to pathway *j*. The extracellular state was

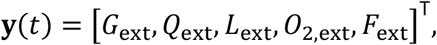

representing extracellular glucose, glutamine, lactate, oxygen, and fatty acids, respectively.

Unless stated otherwise, intracellular and extracellular pools were interpreted in mM, maximum pathway capacities in mM s−1, first-order consumption constants in s−1, and enzyme-capacity multipliers and regulatory factors as dimensionless quantities. The model is a phenomenological coarse-graining of central carbon, redox, and lipid metabolism; it is not a genome-scale reconstruction and does not enforce exact carbon, adenine-nucleotide, or redox conservation.

The default intracellular target state was 5.0 mM ATP, 0.30 mM NADH, 2.0 mM lactate, 1.0 mM glucose, 10^−4^ mM ROS, 0.05 mM NADPH, 0.10 mM pyruvate, 0.20 mM glutamine, 0.15 mM free fatty acids, and 2.0 mM triglyceride. The default static environment contained 11.0 mM glucose, 2.0 mM glutamine, 2.0 mM lactate, 0.20 mM oxygen, and 0.40 mM fatty acids.

#### Generic kinetic functions

A single-substrate saturating reaction was represented by a Michaelis-Menten-type rate law,

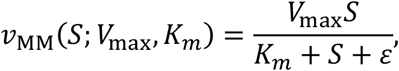

where *ε* = 10^−12^ prevents division by zero. Reversible lumped pathways were represented by independently calculated forward and reverse capacities,

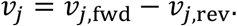

These paired rates are phenomenological pathway-level fluxes and should not be interpreted as elementary reversible enzyme mechanisms.

Adaptive enzyme abundance and experimental perturbations acted primarily on maximum capacity:

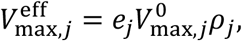

where 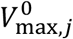 is the baseline capacity, *e*_*j*_ is the enzyme-capacity multiplier, and *ρ*_*j*_ is a multiplicative perturbation override. Under unperturbed initial conditions, *e*_*j*_ = *ρ*_*j*_ = 1. Enzyme adaptation generally scaled *V*_max_ values and did not scale the corresponding *K*_*m*_ values.

Regulatory feedback was applied mainly by modifying the apparent half-saturation constant. For a positive regulatory factor *F*_*j*_(**x**),

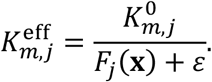

Thus, *F*_*j*_ > 1 increased effective substrate affinity and 0 < *F*_*j*_ < 1 decreased effective affinity. For each metabolite *m*, excess and deficit relative to the target were defined as

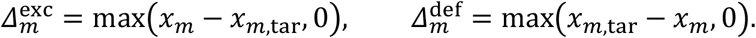

Inhibitory feedback factors had the form

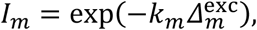

with the exponential argument numerically capped in the implementation. The principal factors were *I*_ATP_, *I*_NADH_, *I*_NADPH_, *I*_Lactate_, *I*_Pyruvate_, *I*_Glutamine_, and *I*_ROS_.

ATP-surplus activation of lipid-storage reactions used a Hill function,

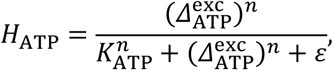

and the bounded surplus multiplier was

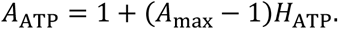

The default values were *K*_ATP_ = 0.5 mM, *n* = 2, and *A*_max_ = 4.

#### Directional transport

Facilitated transport was represented by a saturable carrier term multiplied by a smooth directional gate. For an outside-to-inside gradient,

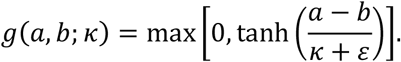

Glucose import was

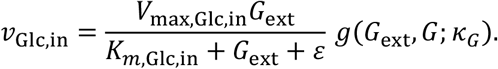

Lactate import and export were computed separately:

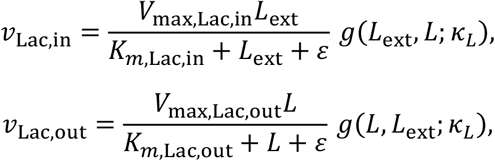

with net cellular lactate transport

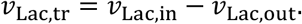

Glutamine import was

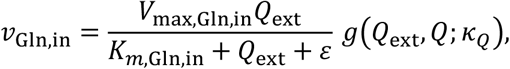

and intracellular glutamine consumption included the first-order term

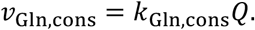

For fatty-acid transport, free extracellular and intracellular fractions were defined as

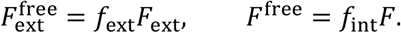

Carrier-mediated import and export were

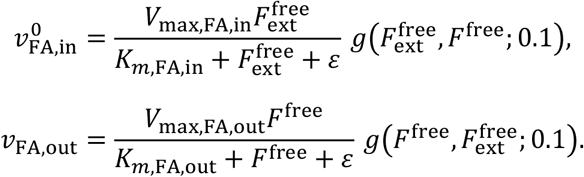

Intracellular fatty-acid and triglyceride excess inhibited inward transport:

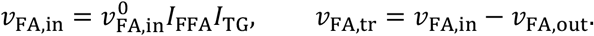

#### Glycolysis and the pentose-phosphate pathway

Cytosolic NAD+ availability was approximated from a fixed total NAD pool,

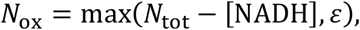

where *N*_tot_ = 1.0 mM by default. Lactate-associated inhibition of glycolysis and the GAPDH redox gate were

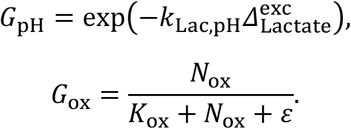

The forward glycolytic feedback multiplier was

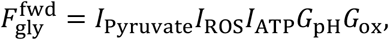

and the reverse multiplier was

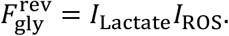

The corresponding rates were

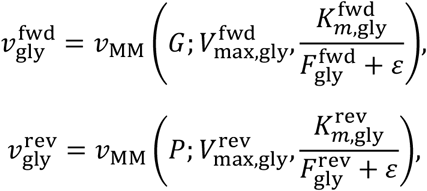

with 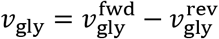.

The pentose-phosphate pathway was activated by elevated ROS and NADPH deficiency. A ROS-dependent boost and NADPH-demand term were defined as

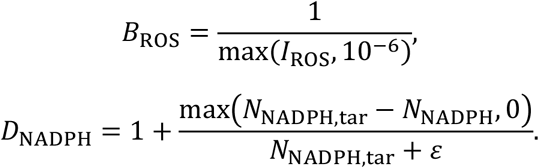

The forward and reverse feedback terms were

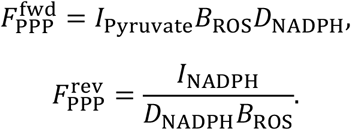

The PPP rates were

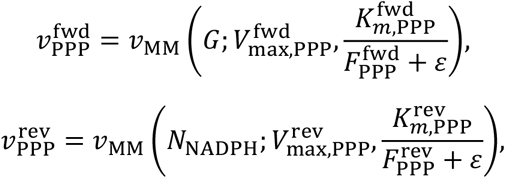

with 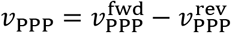.

#### TCA cycle, glutamine anaplerosis, electron transport, and ATPase load

For the pyruvate-fed TCA branch, a common stress term and energy/redox drive were

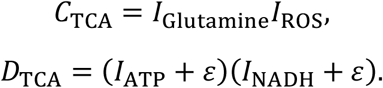

Forward and reverse feedback multipliers were

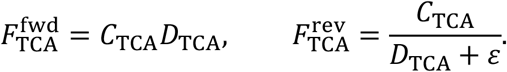

The rates were

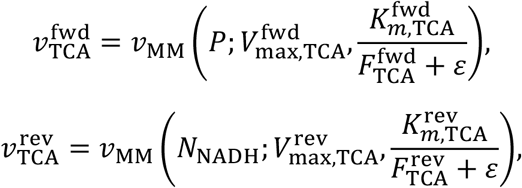

with 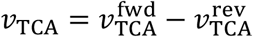.

The glutamine-fed TCA branch used the analogous terms

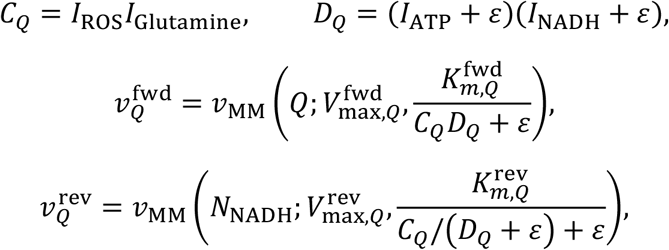

with 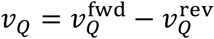.

Electron transport was unidirectional in the model. The oxygen gate was

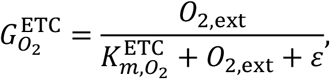

and the ETC feedback term was

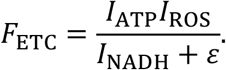

The ETC rate was

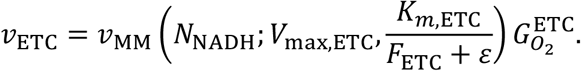

Reverse ETC flux was set to zero. ATP-consuming load was instead represented by an explicit saturable ATPase term,

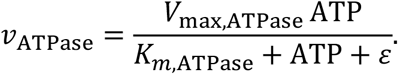

#### Lactate dehydrogenase kinetics

Lactate dehydrogenase was represented as a reversible redox-coupled bi-substrate reaction. The forward reaction was pyruvate plus NADH to lactate plus NAD+,

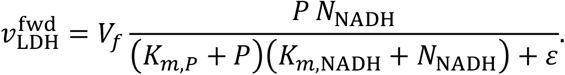

The reverse reaction was lactate plus NAD+ to pyruvate plus NADH,

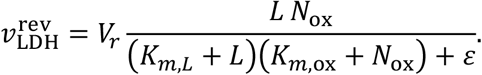

The reverse capacity was derived from the forward capacity using a Haldane-type constraint,

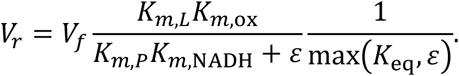

The net LDH rate was

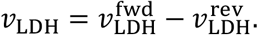

The Haldane-constrained formulation was enabled in the current implementation. The alternative legacy LDH formulation, in which forward and reverse capacities were independently specified and oxygen-gated, was retained only for backward compatibility and was not used in the principal model configuration.

#### Lipid metabolism

Fatty-acid and triglyceride feedback factors were

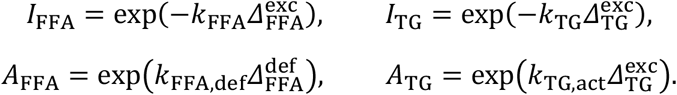

The de novo lipogenesis feedback term was

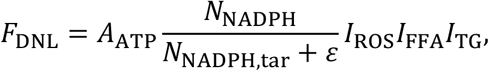

and the DNL rate was

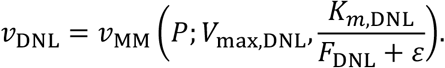

Esterification was

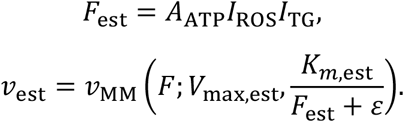

Lipolysis was

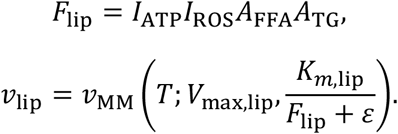

A malonyl-associated inhibitory factor and oxygen gate were

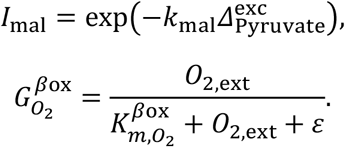

The β-oxidation feedback and rate were

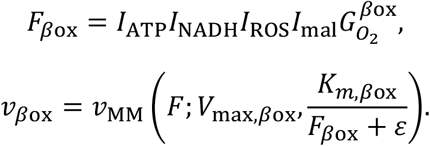

Reverse β-oxidation was neglected at the modeled scale.

#### ROS production and detoxification

ROS production was coupled phenomenologically to positive ETC and β-oxidation fluxes,

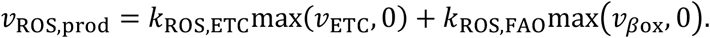

NADPH availability gated ROS detoxification,

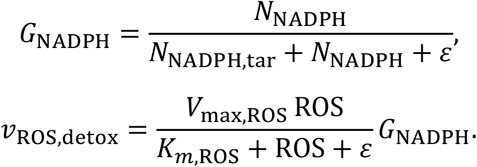

#### Metabolite balance equations

The ten intracellular variables were updated from lumped production, consumption, transport, and maintenance terms. The implemented balance equations were

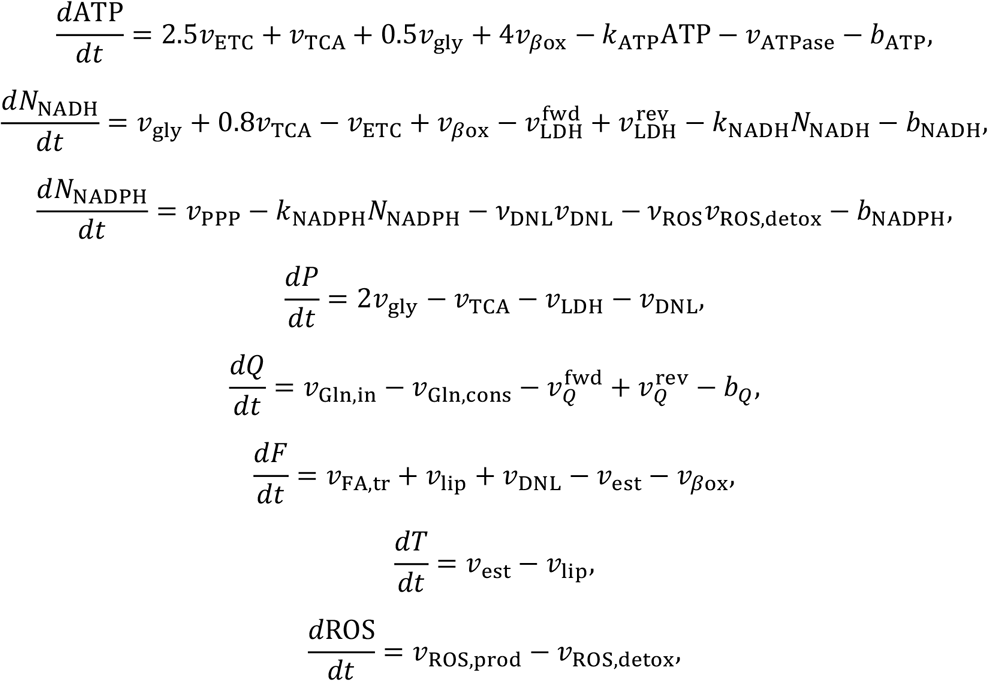

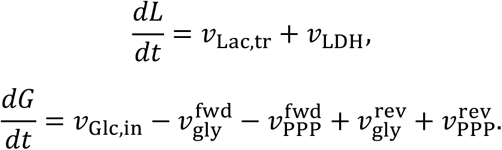

Here, *b*_*m*_ denotes a concentration-independent basal drain and *k*_*m*_*x*_*m*_ denotes a first-order drain. The coefficients 2.5, 0.8, 0.5, and 4 are model-scale yield coefficients rather than exact biochemical stoichiometries.

#### Numerical integration and positivity

The model used an outer time step of *Δt* = 0.1 s. Each outer step was divided into five internal substeps,

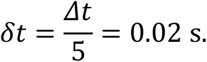

Under the default posdef integrator, each intracellular pool was advanced by explicit Euler integration in linear concentration space,

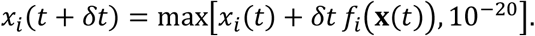

The positive concentrations were stored internally as logarithms for state serialization and assignment, but the default update itself was not a log-space integration. A legacy multiplicative log-Euler update remained available through the MET_MODEL_INTEGRATOR=legacy environment setting but was not the default.

### Adaptive enzyme-capacity control

#### Enzyme representation and bounds

Enzyme activities were represented as dimensionless pathway-capacity multipliers. The model therefore represents adaptive regulation at the level of effective pathway capacity and does not explicitly simulate promoter state, mRNA abundance, translation kinetics, individual protein turnover, allosteric binding, or post-translational modification states.

All enzyme multipliers were initialized at *e*_*j*_ = 1. The default bounds were

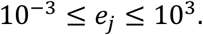

When the optional MCT restriction was enabled, the MCT-associated multiplier was limited to

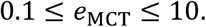

#### Greedy one-step controller

At each outer model step, the controller evaluated a fractional increase and decrease in every enzyme multiplier. For enzyme *j*,

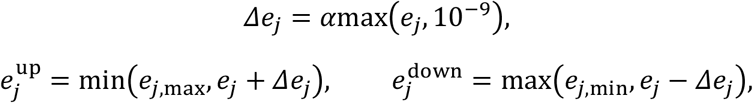

where the default fractional update scale was

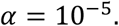

For each candidate, all associated maximum capacities were rebuilt as

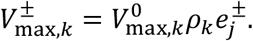

All candidates were evaluated from the same frozen intracellular state, kinetic parameter state, and extracellular environment. Environmental and consumption-rate noise was temporarily disabled during branch evaluation so that the upward and downward alternatives were compared under identical deterministic conditions.

The default controller predicted one outer step ahead. More generally, for a look-ahead length *K*, the candidate state was generated by iterating the metabolic dynamics using a mini-step *Δt*/*K*; in the current default configuration, *K* = 1.

The global control objective was the unweighted relative *L*_1_ deviation from the target state,

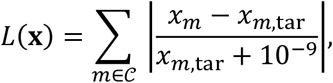

where the default controlled set *C* contained all ten intracellular variables. If local enzyme control was enabled, each enzyme was scored against a pathway-specific subset of substrates, products, and energetic variables; an empty local set fell back to the global control set.

For each enzyme, the candidate with the lowest predicted loss among the three alternatives (decrease, no change, increase) was selected:

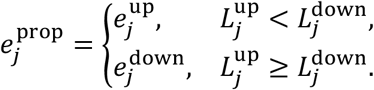

All proposed enzyme values were then committed synchronously. When two or more candidates have losses within a near-tie tolerance of the best, the controller selects the no-change action. This tie-breaking rule prevents slow enzyme drift near a metabolic steady state.

#### Enzyme synthesis, degradation, and maintenance cost

Energetic costs were calculated after the controller selected the proposed enzyme moves. One unit of enzyme multiplier was mapped to an effective enzyme concentration

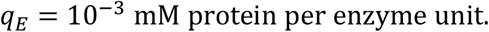

For enzyme *j*, let *a*_*j*_ be the representative protein length in amino acids, *n*_*j*_ the representative transcript length in nucleotides, *s*_j_ the number of phosphorylation sites, and 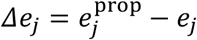. The synthesis cost for *Δe*_*j*_ > 0 was

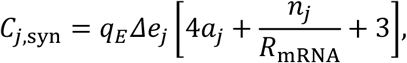

where translation was assigned 4 ATP equivalents per amino acid, transcription 1 ATP equivalent per nucleotide amortized over *R*_mRNA_ = 10^4^ proteins per transcript, and initiation/termination 3 ATP equivalents per synthesized protein.

For *Δe*_*j*_ < 0, degradation cost was

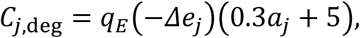

representing 0.3 ATP equivalents per amino acid plus a 5-ATP ubiquitination overhead. The phosphorylation-maintenance cost applied to the proposed extant pool was

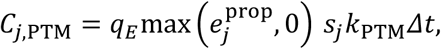

where *k*_PTM_ = 10^−3^ s^−1^ per site and each cycling event cost one ATP equivalent. The total cost was

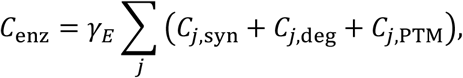

where *γ*_*E*_ is the global enzyme-energy-cost scale. The principal noise-energy sweep used *γ*_*E*_ ∈ {1,10,100}; the run-level value was retained in the output metadata.

When the proposed total cost exceeded the available ATP pool, all proposed enzyme changes were reduced by the common feasibility factor

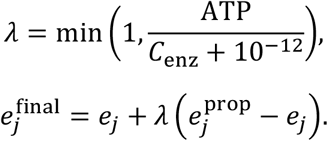

The cost was recomputed for the scaled moves and subtracted directly from the intracellular ATP pool before metabolic integration. Thus, enzyme cost acted as an ATP-resource constraint and debit; it was not included as a penalty in the controller loss *L*.

### Extracellular environment, perturbations, and stochasticity

#### Static and perfused environments

The default environment mode was static: extracellular concentrations remained fixed except when explicitly perturbed. The code also supports a perfused-batch extracellular compartment. For extracellular metabolite *m*, the continuous equation is

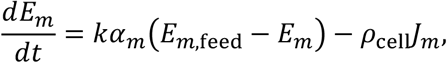

where *k* is the perfusion rate, *α*_*m*_ is a metabolite-specific exchange multiplier, *ρ*_cell_ is a cell-to-environment volume scalar, and *J*_*m*_ is the cell-side uptake term constructed from the most recent intracellular fluxes. A positive *J*_*m*_ removes metabolite from the extracellular compartment.

The perfusion relaxation was integrated semi-implicitly:

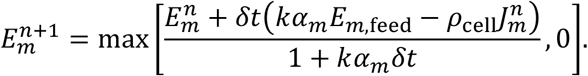

Extracellular and intracellular updates were interleaved at every internal substep. Unless a perfused-batch cohort was explicitly selected, the static environment was used.

#### Perturbation operators

Three perturbation classes were used in the exhaustive escape-attractor search.

For an environmental perturbation of strength *s*, the selected extracellular value was rescaled from its baseline value,

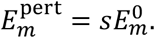

For pathway inhibition, the targeted kinetic capacity was multiplied by

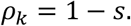

For increased consumption, the selected consumption or production coefficient was multiplied by

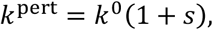

with non-negativity enforcement and an implementation cap of 100 times the pre-perturbation value.

A metabolite-clamp interface exists in helper code, but intracellular clamps were not included in the exhaustive candidate generator and are not consumed by the current state-update equations. They were therefore not treated as an active perturbation class in the present Methods description.

#### Multiplicative noise

Stochasticity could be applied to the environment, to consumption/production coefficients, or to both. Each affected quantity was multiplied by an independent mean-preserving log-normal factor,

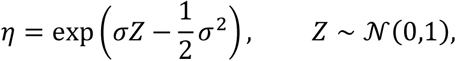

so that

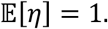

A new multiplier was sampled for each affected variable at each internal substep. The noisy environment or coefficient dictionary was temporary and was restored after the substep, so noise was an instantaneous multiplicative perturbation rather than a persistent random walk. The total NAD pool and LDH equilibrium constant were excluded from consumption/production noise.

The principal factorial search comprised a no-noise baseline and environmental-noise conditions *σ* ∈ {1,2,3}, crossed with enzyme-energy-cost scales *γ*_*E*_ ∈ {1,10,100}. Random seeds were specified per run and retained in the search metadata.

#### Baseline equilibration

Before perturbation assays, each model instance was advanced until either a convergence criterion was satisfied or a maximum number of steps was reached. Convergence was tested every *N*_*c*_ = 100 outer steps by comparing the current metabolite state with the state at the preceding convergence check:

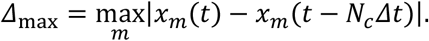

The baseline was classified as converged when

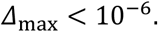

The maximum baseline duration was 500,000 outer steps. With *Δt* = 0.1 s, this corresponded to a maximum of 50,000 s. The convergence test included intracellular metabolites but not enzyme multipliers. The shared baseline-equilibration implementation was used by escape-attractor search, habituation assays, and ischemia-reperfusion workflows, and for baseline initialization wherever those pipelines invoked the canonical equilibration utility.

### Parameter-set generation, admissibility filtering, and selection

#### Parameter sampling

Candidate kinetic parameter sets were generated by multiplicative perturbation of a baseline parameter vector. The baseline was obtained from the default model unless an explicit parameter file or custom parameter dictionary was supplied.

The default sampling method was Latin hypercube sampling in log_10_ parameter space. For parameter *θ*_*k*_ with baseline value 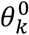,

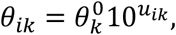

where

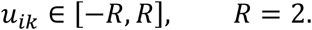

Thus, before repair constraints, each sampled parameter could range from 10^−2^ to 10^2^ times its baseline value. The implementation also supported local Gaussian multiplicative sampling and independent log-uniform sampling, but Latin hypercube sampling was the default.

The default parameter-search seed was 424200 unless overridden. Each Latin-hypercube batch was generated from child seed sequences to provide reproducible but decorrelated batches.

#### Structural repair constraints

Each sampled parameter set was passed through a deterministic structural-repair procedure:

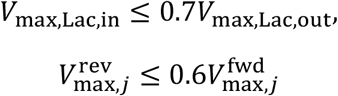

for glycolysis, PPP, TCA, ETC, and the glutamine-fed TCA branch,

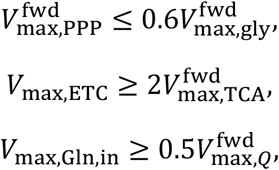

and

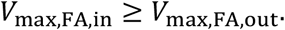

These constraints imposed broad pathway-capacity priors and prevented extreme sampled combinations from generating obviously degenerate pathway wiring. They did not constitute a comprehensive mechanistic prior.

#### Search simulation and admissibility envelope

Unless overridden, each repaired candidate was simulated with *Δt* = 0.1 s for 250,000 s. Only the final 10,000 outer time points were retained for scoring. At the default time step, this tail represented the final 1,000 s rather than 10,000 s.

Candidate simulations were terminated early if any tracked metabolite left a prespecified admissibility envelope. For metabolites without an absolute bound, the effective target was

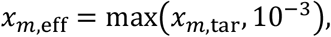

and the admissible interval was

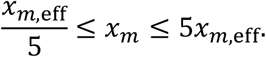

ROS used the absolute interval

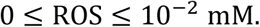

The admissibility test was applied after every outer step, beginning with the first step. Runs that violated the envelope were excluded from the main search log and did not count toward the requested number of retained parameter sets. Sampling continued until the requested number of admissible runs was obtained; the default target was 1,000 admissible runs. The envelope is an operational numerical/physiological filter and should not be interpreted as a validated biological cell-death model.

#### Scoring functions

Metabolic stability was quantified over the retained terminal window. If the window comprised indices *t* = *i*_0_, …, *i*_1_ − 1, the stability loss was

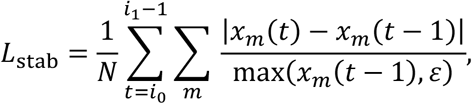

where *N* is the number of evaluated metabolite-step pairs. Smaller values indicated less step-to-step variation.

An analogous enzyme-stability loss was calculated over the same shared terminal interval when adaptive enzyme control was enabled.

Target loss was the summed absolute final log_10_ deviation from the target state,

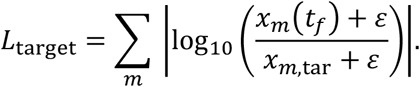

The combined loss was

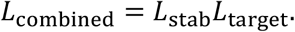

The search objective was selected by configuration. The default objective mode was metabolic stability, so parameter sets were ranked by *L*_stab_ unless target or combined mode was explicitly requested. Search records retained all three losses, enzyme-stability loss, parameter values, run identity, attempted and excluded counts, simulation length, noise configuration, and enzyme-control settings.

### Baseline profiling of selected parameter sets

Baseline profiling summarized long-duration behavior independently of acute perturbation assays. By default, the profiler selected the ten admissible search records with the smallest target loss, not necessarily the smallest stability loss. Each selected parameter set was simulated under baseline conditions for 500,000 s unless a different profiling duration was supplied.

The profiler retained a rolling terminal window whose length was inherited from the search record, normally 10,000 time points, corresponding to 1,000 s at *Δt* = 0.1 s. It calculated terminal values and tail means for metabolite states, forward and reverse pathway rates, net pathway rates, enzyme multipliers, first-order and basal consumption components, and ATP-equivalent enzyme-cost components. It also exported sampled trajectories and checkpoint summaries.

The run-level enzyme-step size, energy-cost scale, noise parameters, and admissibility settings were propagated from the search record or run metadata where available. Because the profiler currently contains legacy fallback values for missing enzyme-control metadata, complete run metadata are required for reproducible profiling.

### Ischemia-reperfusion assay

After baseline equilibration, ischemia was imposed by setting

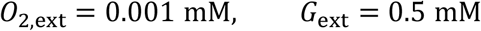

for 10,000 s. Reperfusion was initiated by abruptly restoring extracellular oxygen and glucose to their pre-ischemic baseline values and continuing the simulation for 40,000 s in the principal single-experiment runner.

The assay recorded the ATP nadir during ischemia, the maximum ROS concentration during ischemia, the maximum ROS concentration during reperfusion, and the ROS peak ratio

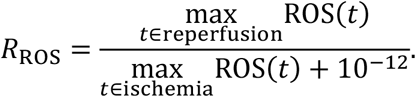

A parameter set was operationally classified as displaying an ischemia-reperfusion-like ROS pattern when

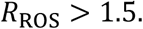

This classification evaluates the model’s phenomenological energetic and oxidative response. The model does not explicitly represent succinate accumulation, reverse electron transport, permeability transition, mitochondrial membrane damage, or antioxidant-enzyme damage.

### Escape-attractor search

Escape-attractor search identified perturbation pulses that produced the largest persistent displacement from the current baseline state after perturbation removal and recovery. Each selected parameter set was first equilibrated using the canonical baseline procedure.

The exhaustive candidate generator tested: (i) inhibition of every *V*_max_-like kinetic parameter; (ii) increases in first-order or basal consumption coefficients and the principal ETC-associated ROS-production coefficient; and (iii) rescaling of each extracellular environmental variable. The default strengths were

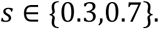

The default search contained two sequential rounds. In each round, a candidate perturbation was applied for 100,000 outer steps and then reverted. The model was subsequently advanced for a recovery interval of 400,000 outer steps. At *Δt* = 0.1 s, these intervals corresponded to 10,000 s and 40,000 s, respectively.

Candidate metabolic displacement was measured at the end of recovery using the raw, unnormalized *L*_1_ distance

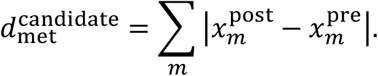

Candidate enzyme displacement was

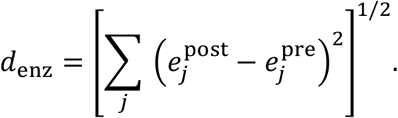

The selected perturbation was then reapplied to the working model, reverted after the perturbation interval, and followed by the recovery interval. The committed roundwise displacement was reported using Euclidean metabolic distance,

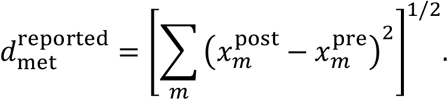

After each committed round, the model was optionally re-equilibrated for up to 200,000 steps using an absolute tolerance of 10^−7^ before the next search round. Outputs included the selected perturbation sequence, complete per-candidate rankings, metabolic and enzyme displacements, perturbation timing, strengths, parameter identity, and sampled simulation traces.

### Habituation protocol

#### Stimulus selection

The training stimulus P and probe stimulus Q were selected from the escape-attractor search (EAS) round-1 screen, which evaluates all perturbation channels at two strengths from the same equilibrated attractor. The screen selects a matched pair per parameter set (rank) according to five criteria: (1) responsive – naive peak displacement ≥ 5× the noise floor (10th percentile of all peaks); (2) recovers – fractional recovery 1 – d_residual / d_peak ≥ 0.80; (3) has headroom – peak < 50 (rejects saturated responses); (4) different channels – P and Q act through different metabolites or reactions; and (5) matched amplitude – max(P_peak, Q_peak) / min(P_peak, Q_peak) ≤ 1.25.

| Role | Perturbation | Target | Clamp value | Effective env. |
| --- | --- | --- | --- | --- |
| P (train) | change_env, strength 0.3 | extracellular fatty acid | 0.4 mM | 0.12 mM |
| Q (probe) | change_env, strength 0.3 | extracellular lactate | 2.0 mM | 0.6 mM |

Naive reference peaks: P_peak_ = 7.63, Q_peak_ = 6.91 (L1 displacement), amplitude ratio 1.10. P enters through the FA_Import transporter (*K*_m_ = 0.10 mM); Q enters through MCT (*K*_m_ = 1.0 mM). The two transport pathways are independent.

#### Protocol structure

Each cell runs the following protocol after a 10^6^ s baseline settle:

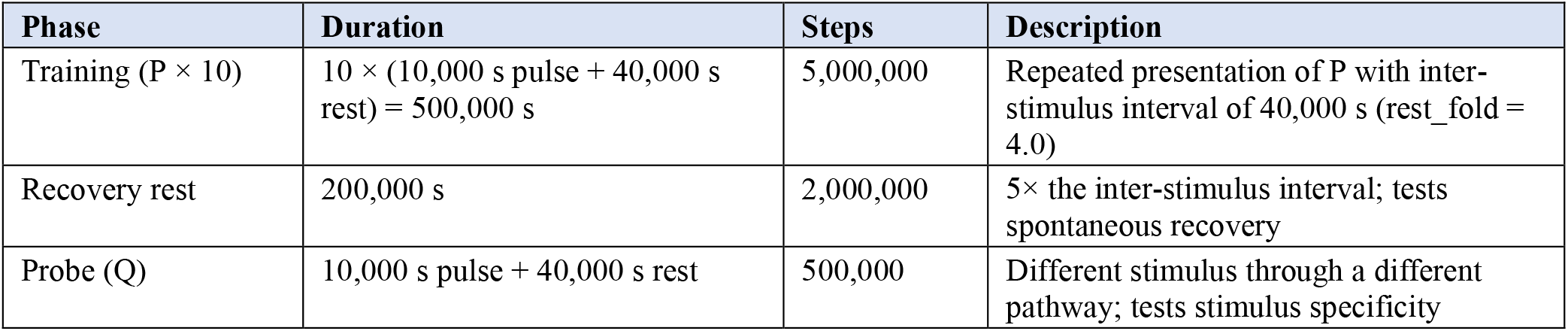
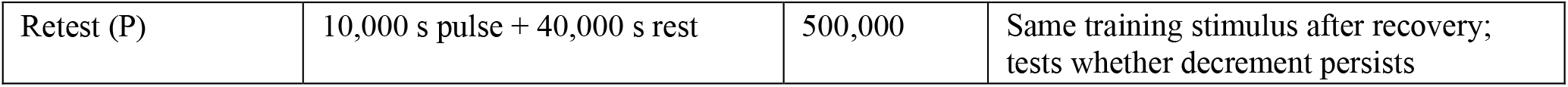

Each cycle’s displacement is measured against the metabolic state recorded at the start of that cycle, so the reported values are evoked responses with no standing offset mixed in. The displacement metric is L1 (sum of absolute relative deviations across 10 metabolites: ATP, FFA, glucose, glutamine, lactate, NADH, NADPH, pyruvate, ROS, and triglyceride). The residual displacement is measured against the baseline equilibrium state.

#### Experimental arms

Each protocol runs on two arms in parallel. The adaptive arm runs the bang-bang enzyme controller (α = 10^−5^ per step, energy_cost_scale = 100) with the transport desensitisation mechanism enabled at the specified *τ*_hab_ and *τ*_rec_. The frozen arm fixes enzymes at their settled levels with no controller adjustments and no desensitisation, serving as the no-plasticity control.

An additional off cell (adaptive arm, no desensitisation) serves as the baseline for comparison: any response decrement in the off cell comes from the controller dynamics alone, not from the desensitisation mechanism.

#### Parameter sweep

The desensitisation mechanism has two free parameters: *τ*_hab_ (accumulation timescale) and *τ*_rec_ (recovery timescale). We swept a 3 × 3 grid:

*τ*_hab_ ∈ {3,000, 10,000, 30,000} s

*τ*_rec_ ∈ {30,000, 100,000, 300,000} s

plus an off cell (no desensitisation), for a total of 10 cells per host. The sweep was run on three hosts: none_ec100_rank8 (the primary host, energy_cost_scale = 100), none_ec100_rank3, and none_ec100_rank5.

#### Measurements

For each stimulus presentation, the following quantities were recorded: (1) peak displacement – the maximum L1 distance from the pre-cycle state during the pulse, serving as the primary habituation readout; (2) rebound displacement – the largest excursion during the rest phase after the clamp is released (often larger than the peak displacement, since the release itself is a perturbation); (3) mean displacement – the average L1 distance over the stimulus window; (4) residual displacement – the L1 distance from the baseline equilibrium at the end of the rest period; and (5) full metabolite and enzyme traces – all 10 metabolite concentrations and 15 enzyme multipliers sampled every 100 steps (10 s).

#### Habituation criteria

Habituation was assessed against the Thompson & Spencer (1966) criteria. Response decrement was measured as a monotonic decline in peak displacement over the 10 training cycles, quantified as 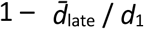, where 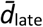 is the mean of the last three training peaks and *d*_1_ is the first. Spontaneous recovery was measured as the retest response *d*_retest_ after the 200,000 s recovery rest, quantified as *d*_retest_ / *d*_1_ (recovery fraction). Stimulus specificity was assessed by comparing the probe response *d*_probe_ to the off cell’s probe response. Habituation to P should not transfer to Q.

### Alternative mechanisms tested

In addition to the transport desensitisation mechanism described above, the following controller variants and plasticity mechanisms were tested on the same protocol and parameter set. Their outcomes are reported in the Results.

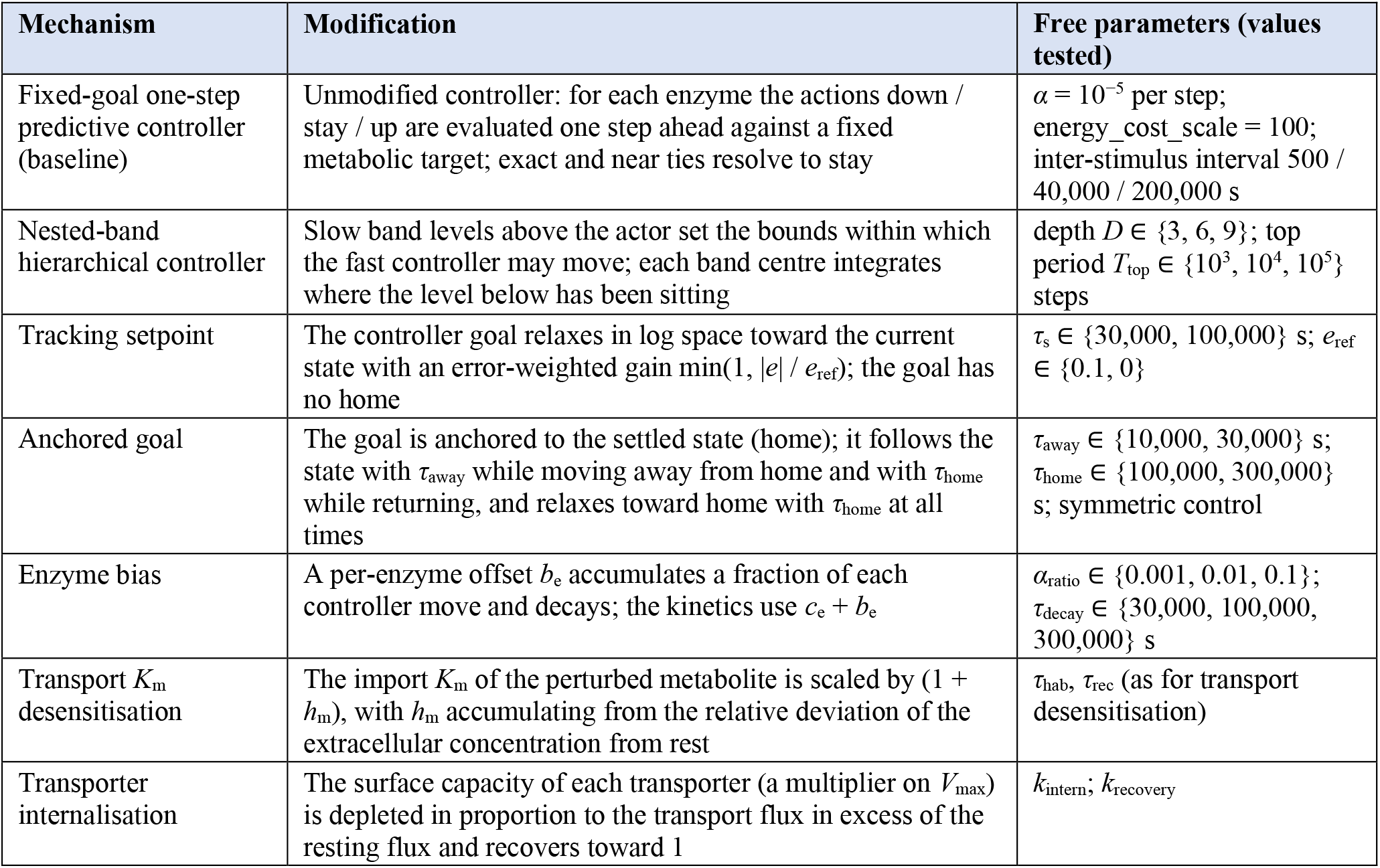

## Data Availability

All data generated during the current study are included in this published article. Simulation outputs are reproducible from the code and parameters described in the Methods section.

## Code Availability

The underlying code for this study will be made available in a public repository upon publication.

## References

1. Nijhout, H.F., J.A. Best, and M.C. Reed, Systems biology of robustness and homeostatic mechanisms. WIREs Systems Biology and Medicine, 2019. 11(3): p. e1440.

2. Watson, E., L.S. Yilmaz, and A.J.M. Walhout, Understanding Metabolic Regulation at a Systems Level: Metabolite Sensing, Mathematical Predictions, and Model Organisms. Annual Review of Genetics, 2015. 49(Volume 49, 2015): p. 553–575.

3. Baluška, F. and M. Levin, On Having No Head: Cognition throughout Biological Systems. Front Psychol, 2016. 7: p. 902.

4. Kaygisiz, K. and R.V. Ulijn, Can Molecular Systems Learn? ChemSystemsChem, 2024. 7: p. e202400075.

5. Kukushkin, N.V., et al., The massed-spaced learning effect in non-neural human cells. Nat Commun, 2024. 15(1): p. 9635.

6. Levin, M., The Multiscale Wisdom of the Body: Collective Intelligence as a Tractable Interface for Next-Generation Biomedicine. Bioessays, 2024: p. e202400196.

7. Mathews, J., et al., Cellular signaling pathways as plastic, proto-cognitive systems: Implications for biomedicine. Patterns (N Y), 2023. 4(5): p. 100737.

8. Keresztes, D., et al., Cancer drug resistance as learning of signaling networks. Biomed Pharmacother, 2025. 183: p. 117880.

9. Veres, T., et al., Cellular forgetting, desensitisation, stress and ageing in signalling networks. When do cells refuse to learn more? Cell Mol Life Sci, 2024. 81(1): p. 97.

10. Csermely, P., et al., Learning of Signaling Networks: Molecular Mechanisms. Trends Biochem Sci, 2020. 45(4): p. 284–294.

11. Chandel, N.S., Metabolism of proliferating cells. Cold Spring Harbor Perspectives in Biology, 2021. 13(10): p. a040618.

12. Hensley, C.T., et al., Metabolic heterogeneity in human lung tumors. Cell, 2016. 164(4): p. 681–694.

13. Biswas, S., W. Clawson, and M. Levin, Learning in transcriptional network models: computational discovery of pathway-level memory and effective interventions. International journal of molecular sciences, 2022. 24(1): p. 285.

14. Biswas, S., et al., Gene regulatory networks exhibit several kinds of memory: Quantification of memory in biological and random transcriptional networks. Iscience, 2021. 24(3).

15. Pigozzi, F., A. Goldstein, and M. Levin, Associative conditioning in gene regulatory network models increases integrative causal emergence. Communications Biology, 2025. 8(1): p. 1027.

16. Orth, J.D., I. Thiele, and B.Ø. Palsson, What is flux balance analysis? Nature biotechnology, 2010. 28(3): p. 245–248.

17. Smallbone, K., et al., A model of yeast glycolysis based on a consistent kinetic characterisation of all its enzymes. FEBS letters, 2013. 587(17): p. 2832–2841.

18. Villaverde, A.F., et al., A protocol for dynamic model calibration. Briefings in bioinformatics, 2022. 23(1): p. bbab387.

19. Schwanhäusser, B., et al., Global quantification of mammalian gene expression control. Nature, 2011. 473(7347): p. 337–342.

20. Davidson, Shawn M., et al., Environment Impacts the Metabolic Dependencies of Ras-Driven Non-Small Cell Lung Cancer. Cell Metabolism, 2016. 23(3): p. 517–528.

21. Chen, M., et al., Influence of Progressive Tumor Growth on Glutamine Metabolism in Skeletal Muscle and Kidney. Annals of Surgery, 1993. 217(6): p. 655–667.

22. Wegner, A., et al., How metabolites modulate metabolic flux. Current Opinion in Biotechnology, 2015. 34: p. 16–22.

